# Single-cell atlas of tumor clonal evolution in liver cancer

**DOI:** 10.1101/2020.08.18.254748

**Authors:** Lichun Ma, Limin Wang, Ching-Wen Chang, Sophia Heinrich, Dana Dominguez, Marshonna Forgues, Julián Candia, Maria O. Hernandez, Michael Kelly, Yongmei Zhao, Bao Tran, Jonathan M. Hernandez, Jeremy L. Davis, David E. Kleiner, Bradford J. Wood, Tim F. Greten, Xin Wei Wang

## Abstract

Tumor evolution is a key feature of tumorigenesis and plays a pivotal role in driving intratumor heterogeneity, treatment failure and patients’ prognosis. Here we performed single-cell transcriptome profiling of 46 primary liver cancers from 37 patients enrolled for interventional studies. We surveyed the landscape of ~57,000 malignant and non-malignant cells and determined tumor cell clonality by developing a machine learning-based consensus clustering method. We found evidence of tumor cell branching evolution using hierarchical clustering, RNA velocity as well as reverse graph embedding methods. Interestingly, an increasing tumor cell clonality was tightly linked to patients’ prognosis, accompanied by a polarized immune cell landscape. We identified osteopontin as a key player for tumor cell evolution and microenvironmental reprogramming. Our study offers insight into the collective behavior of tumor cell communities in liver cancer as well as potential drivers for tumor evolution in response to therapy.

## INTRODUCTION

Tumorigenesis is a consequence of an evolutionary process by which somatic cells acquire genetic and epigenetic alterations when exposed over time to extrinsic or intrinsic factors such as adverse tumor microenvironment (TME) (Maley et al., 2017). It is known that tumor cells within a solid lesion exist as a social community where each tumor cell can interact with one another, as well as their TME that includes extracellular matrix, tumor vasculature and immune cells. This TME provides a setting in which tumor cells sense and adapt to environmental cues such as available nutrients, oxygen, as well as toxic agents such as chemotherapeutics for its survival and growth (Junttila and de Sauvage, 2013). As such, a tumor lesion is under the constant pressure of Darwinian evolutionary selection reminiscent to the ecology of collective behavior of the animal kingdom (Gordon, 2014). Consequently, such an intrinsic ‘survival of the fittest’ trait, found in most if not all malignant solid tumors, can result in vast heterogeneity within tumor cell populations known as intratumor heterogeneity (ITH) (Maley et al., 2017), a feature universally linked to tumor aggressiveness (Andor et al., 2016). Evidence of ITH has been observed in multiple tumor types including brain, breast, colon, head and neck, liver, lung and skin cancer by both tumor bulk and single cell analyses (Azizi et al., 2018; Gerlinger et al., 2012; Li et al., 2017; Ma et al., 2019; Puram et al., 2017; Tirosh et al., 2016a; Tirosh et al., 2016b; Venteicher et al., 2017). In the same way that biodiversity within a defined ecosystem promotes fitness and survival of an organism, ITH has a similar role in sustaining tumor survival. Thus, presence of ITH creates unique challenges for targeted therapy, leading to drug resistance, treatment failure and poor outcomes (Khatib et al., 2020). This begs the questions of what are cellular factors driving the collective behaviors of a tumor cell community? Are there intrinsic drivers regulating tumor cell evolution, similar to the proposed model by referring ITH as a reflection of the inherent biology of a given tumor type (Iacobuzio-Donahue et al., 2020; Khatib et al., 2020)? Understanding the biodiversity and clonality of solid cancers on a cellular or lesion-specific basis may provide conceptual knowledge about carcinogenesis with practical implications for diagnosis and treatment.

Primary liver cancer is among the top five deadliest cancers in the world, of which the most common types are hepatocellular carcinoma (HCC) and intrahepatic cholangiocarcinoma (iCCA) (Bray et al., 2018). Both tumor types have been shown to have extensive intertumor and intratumor heterogeneity (Boyault et al., 2007; Chaisaingmongkol et al., 2017; Jusakul et al., 2017; Lee et al., 2004; TheCancerGenomeAtlasResearchNetwork, 2017). This is unsurprising given the presence of multiple and complex etiological factors, extensive genomic complexity and underling tumor biology in HCC and iCCA (Wang and Thorgeirsson, 2014). While most HCC and iCCA are refractory to systemic therapeutics that target tumor cells (Worns and Galle, 2014), some HCC show remarkably durable responses to immune checkpoint inhibitors either alone or in combination with ablation or anti-VEGF (Duffy et al., 2017; Finn et al., 2020). A recent study using single cell transcriptome analysis offered a mechanistic explanation for differential treatment responses, and provided a rationale for use of a combination therapy of immune checkpoint blockade and anti-vascular treatment to improve therapeutic efficacies (Finn et al., 2020; Ma et al., 2019). Thus, the single cell transcriptome analysis provides sufficient resolution to faithfully determine tumor cell communities. Here, using single cell transcriptome analysis, we aimed to delineate tumor cell evolution by examining 46 HCC and iCCA biospecimens from 37 patients enrolled in various treatment protocols at the NIH Clinical Center, with 16 biospecimens collected before and after treatment from 7 patients. We validated our findings in bulk transcriptomic data from 488 patients with HCC and 277 patients with iCCA. Our study provides rich resources and valuable insights into the understanding of a complex liver cancer ecosystem.

## RESULTS

### Single-cell atlas of tumor ecosystem in liver cancer

To study tumor evolution, we performed single-cell RNA sequencing (scRNA-seq) of liver tumors from 37 liver cancer patients participating in interventional clinical trials at the NIH Clinical Center (Figure 1A and Table S1). In a subset of seven patients, liver biopsies were available from multiple time points, including both before and after treatment. We obtained single-cell transcriptomic profiles of 56,721 single cells (Figure 1B). Epithelial marker genes were highly expressed in some of the cells, suggesting potential tumoral origins (Figure 1C). For a representative patient (H34) with three biopsies collected, we found a relatively similar tumor ecosystem, with similar cell types such as T cells, B cells, cancer-associated fibroblasts (CAFs), tumor-associated macrophages (TAMs), tumor-associated endothelial cells (TECs) and even epithelial cells demonstrating similar populations between biopsies collected at different times, revealing relatively stable tumor stromal compositions over time (Figures 1D and S1A). We further confidently identified 17,164 malignant cells using a method of inferring genome-wide chromosomal copy number variations (CNVs) from scRNA-seq data as previously described (Ma et al., 2019) (Figures S1B-S1D). We found malignant cells formed patient-specific clusters (Figure 1E, top panel), consistent with previous studies (Ma et al., 2019; Neftel et al., 2019; Puram et al., 2017; Tirosh et al., 2016a). Again, a heterogeneous population was observed within each cluster, indicating the presence of both intertumor and intratumor heterogeneity with a greater heterogeneity between tumors than within a tumor. This was also validated by measuring the pairwise correlation of single cells, where the single cells within a tumor are more strongly correlated with each other than with those between tumors (Figure 1E, bottom panel). In contrast to the vast heterogeneity of malignant cells, non-malignant cells mainly grouped according to cell types, as annotated according to known cell lineage specific marker genes unique to T cells, B cells, CAFs, TAMs, TECs, hepatocytes and cholangiocytes (Figure 1F). We confirmed the histology of liver tumors in our study by histopathology analysis (Figure 1G).

**Figure 1.**
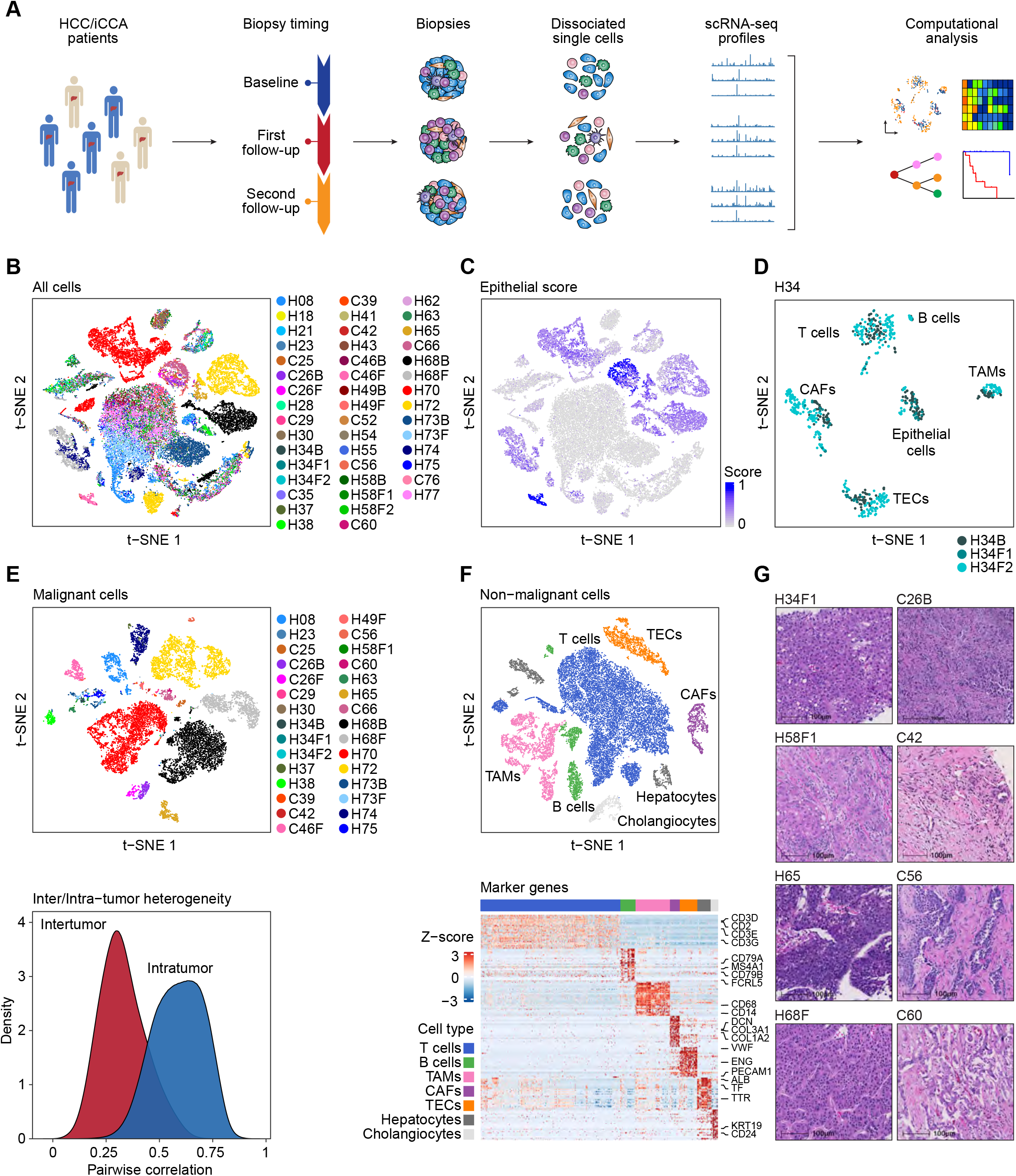
Single-cell transcriptomic profiling of primary liver tumor. (A) Workflow of liver tumor biopsy collection, processing, sequencing, and computational analysis. (B) t-SNE plot of 56,721 single-cells from 46 liver tumor samples (indicated by colors). Case ID was named according to histological subtypes of liver cancer as well as biopsy timing. H, HCC; C, iCCA; B, baseline (the first biopsy); F, follow-up biopsy after treatment. (C) t-SNE plot of single-cells in (B) colored by epithelial score. Epithelial score was determined by the average expression of epithelial marker genes. (D) t-SNE plot of a representative case (H34) with biopsies collected at baseline and follow-up study during treatment. (E) t-SNE plot (top) and tumor heterogeneity (bottom) of malignant cells from 30 tumors with > 15 malignant cells in each tumor. (F) t-SNE plot (top) and known lineage-specific marker genes (bottom) of non-malignant cells. Cell types were indicated by colors. (G) Histopathology of the eight representative tumors. Scale bars, 100 μm.

### Tumor cell clonality and tumor branching evolution

We determined the functional clonality of malignant cells within each tumor by developing a consensus clustering algorithm to ensure the stability of clone detection as well as to account for imbalance of the number of malignant cells between tumors (Figures S2A and S2B). With the derived clones, we constructed the tumor branching architecture based on hierarchical clustering method (Figure 2A, top two panels). Bootstrap analysis (Suzuki and Shimodaira, 2006) was applied to measure the confidence of the hierarchical relationship of the clones (Figure S2C, top panel). We found three major branches in the phylogenetic tree and referred the branches using branching index (BI) as BI-A, BI-B, and BI-C. All tumors in BI-A are from HCC cases, while BI-B and BI-C contain tumors from both HCC and iCCA. In all three branches, clones within a tumor tend to be grouped together forming a hierarchical architecture. To further determine clonal relationship within a tumor, we applied two independent methods: reverse graph embedding method (Qiu et al., 2017) and RNA velocity (La Manno et al., 2018), to learn the single-cell trajectory and cell-lineage among malignant cells (Figure 2A, middle panels rows 3-5). Both methods revealed similar tumor cell trajectories, which largely agree with the clonal relationship within each tumor using a hierarchical clustering method (Figure 2A, top panel). To determine potential drivers of tumor branching evolution, we developed a strategy to search for conserved genes of different branches (Figure 2B). We reasoned that a gene that is elevated in tumor cells and expressed ubiquitously at the branch or sample level, is likely to drive the function of the corresponding branch or sample. This strategy revealed SPP1 among the top of the conserved genes in BI-B and BI-C (Figure S2C, bottom panel). Interestingly, SPP1 expression was often enriched in tumor cells at the beginning of a cell lineage as determined by the RNA velocity method (Figure 2A, panel row 6). In general, SPP1 expression was much higher in BI-B/C than BI-A (Figure 2A, bottom panel). Moreover, SPP1 was mainly expressed in malignant cells rather than non-malignant cells (Figure 2C). These results suggest that SPP1 may be a driver of tumor branching evolution.

**Figure 2.**
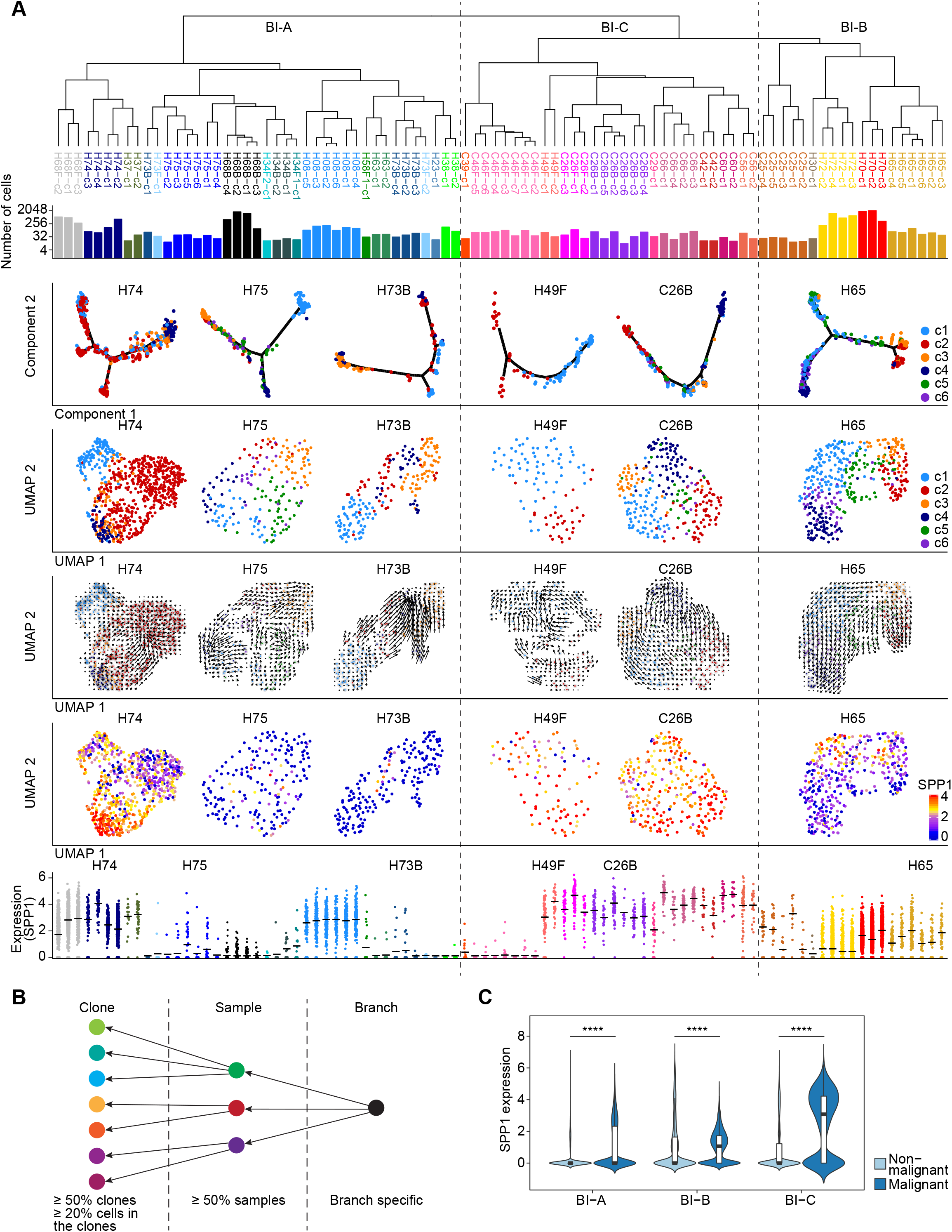
Tumor cell clonality and tumor branching evolution. (A) Tumor phylogenetic tree constructed by hierarchical clustering of all the clones from 30 tumors (indicated by colors) with > 15 malignant cells. BI-A, BI-B and BI-C were defined according to the hierarchical relationship. Panels from top to bottom: phylogenetic tree (top); the number of cells in each clone (row 2); single-cell trajectory (row 3), tumor cell clonality (row 4), RNA velocity-based cell lineage (row 5), and SPP1 expression (row 6) of several representative tumors; SPP1 expression in each subclone of the 30 tumors, with representative tumors labeled on top of the jitter plot (bottom). Line segments in the jitter plot indicate the mean values. (B) Strategy used for searching conserved genes of tumor branches. (C) SPP1 expression in malignant cells and non-malignant cells of different branches. Wilcoxon test was performed to indicate statistical significance. The width of a violin plot represents the density of gene expression values. Box spans the first quartile to the third quartile of the values while segment inside a box indicates the median value. **** p-value < 0.0001.

### Tumor clonality is associated with patient prognosis

We determined the number of clones within a tumor among different branches of the tumor cell phylogenetic tree. Remarkably, we found that there was an increasing number of clones in tumors from BI-A, BI-B, and BI-C (Figure S3A), suggesting that tumors in BI-B and BI-C were more diverse and thus likely more aggressive than tumors in BI-A, a feature described in previous studies (Andor et al., 2016; Kwon et al., 2019; Ma et al., 2019). To further assess tumor aggressiveness, we divided patients into two groups according to the number of clones within a tumor. Strikingly, patients whose tumor had a higher clone number had a much shorter survival than patients with a lower clone number, suggesting a link of tumor clonality and patient prognosis (Figure 3A). In addition, there was a significant trend of overall survival of patients in BI-A, BI-B and BI-C (Figure 3B). Due to a small number of cases in BI-B, we combined BI-B and BI-C as BI-B&C to increase the statistical power. Consistently, patients in BI-B&C had a significantly abbreviated survival than patients in BI-A (Figure 3C). To avoid confounding factors of histological subtypes of HCC and iCCA, we also performed survival analysis of HCC patients separately. Consistently, a significant difference in overall survival was observed in patients between BI-A and BI-B&C (Figure 3D). We did not perform separate analysis for iCCA as they were found only in BI-B&C.

**Figure 3.**
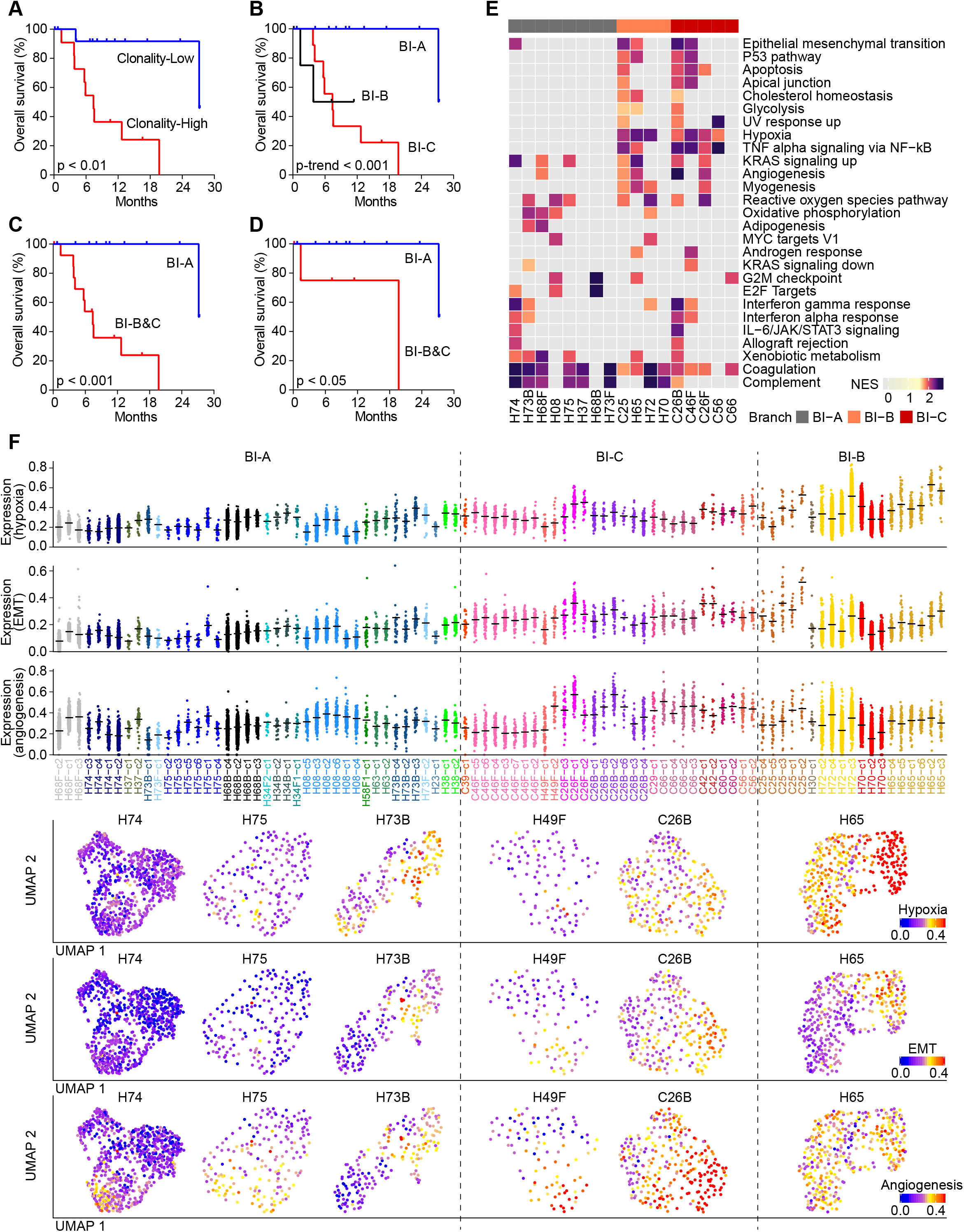
Tumor branching evolution is associated with patient prognosis. (A) Overall survival of all patients with low number of clones (Clonality-Low) and high number of clones (Clonality-High) separated by the median value. (B and C) Overall survival of all the patients from different branches, by using three-group comparison (B) and two-group comparison (C). (D) Overall survival of HCC patients from different branches. (E) The enriched pathways of tumors from BI-A, BI-B and BI-C. Only tumors with more than two clones detected as well as at least one enriched pathway with FDR adjusted q value < 0.05 were involved in the heatmap. NES, normalized enrichment score. (F) Functional analysis of clones within each tumor. Top three panels show the expression of hypoxia, EMT and angiogenesis related genes in each subclone of the tumors from BI-A, BI-B and BI-C. Bottom three panels show the expression of hypoxia, EMT and angiogenesis related genes on the UMAP of several representative tumors. Line segments in the jitter plot indicate the mean values. Log-rank test and log-rank test for trend were preformed to show the statistical significance in (A-D).

To determine signaling pathways enriched in BI-B&C, we carried out gene enrichment analysis of each tumor based on differential expression genes among all the clones within a tumor. We found that tumors in BI-B and BI-C were enriched in tumor aggressiveness-related pathways, such as epithelial–mesenchymal transition (EMT), hypoxia, angiogenesis, TNF alpha signaling as well as glycolysis (Figure 3E). We then examined the tumors from different branches for the aggressiveness-related pathways, such as hypoxia, EMT, and angiogenesis by using the genes from GSEA (Table S2). Noticeably, all the three pathway-related genes were elevated in BI-B and BI-C compared with BI-A (Figure 3F, top three panels). Moreover, the genes were heterogeneously expressed among clones within a tumor, suggesting that different clones may serve different roles in a tumor ecosystem (Figure 3F, bottom three panels). These results are consistent with the survival analysis described above, indicating that patients in BI-B and BI-C tend to gain aggressive tumor features.

Our single cell results described above were based on limited number of cases with varying clinical presentations. To avoid potential confounding factors and to further validate the findings from our single-cell study, we analyzed bulk transcriptomic data from two HCC cohorts of 488 patients (i.e., Liver Cancer Institute, LCI, cohort and TCGA HCC cohort) and two iCCA cohorts of 277 patients (i.e., International Cancer Genomics Consortium, ICGC, cohort and Japan cohort) with >100 cases in each cohort with available survival data. All tumors in the four cohorts were resected specimens. We conducted differential gene expression analysis among malignant cells from BI-B&C and BI-A to generate tumor clonality surrogate gene signatures. The derived signatures from BI-A and BI-B&C were then applied in the four cohorts for survival risk prediction, respectively. We obtained remarkably consistent results from the four cohorts, with a better separation of high-risk and low-risk groups of patients as well as a higher hazard ratio by using the surrogate signature from BI-B&C than the signature from BI-A (Figures S3B and S3C). We also applied tumor clonality surrogate signatures generated from malignant cells of HCC cases only into survival analysis of the four cohorts. Consistent results were obtained (Figures S3B and S3D). Taken together, our results indicate that tumor clonality is associated with patient prognosis, independent of tumor types.

### Polarization of T-cell landscape with tumor branching evolution

We performed t-SNE analysis of T cells, B cells, TAMs, CAFs and TECs derived from three main tumor branches, and found that these cells differed in their transcriptomic profiles between BI-A and BI-B&C, suggesting a potential reprogramming of non-malignant cells by malignant cells (Figure S4A). We focused mainly on T cells for further analysis (Figure S4B; see Methods for details), as tumor-associated T cell subtypes have been extensively characterized by single cell transcriptome analysis (Guo et al., 2018; Zhang et al., 2018; Zhang et al., 2019; Zheng et al., 2017). To confidently determine T cell subtypes, we first extracted CD4^+^ or CD8^+^ T cells by using marker genes of CD4, CD8A or CD8B, respectively. A graph-based clustering method was then applied to define T-cell subtypes according to known marker genes (Figures S4C-S4H), following the approach described in the recent studies (Guo et al., 2018; Zhang et al., 2019; Zheng et al., 2017). We finally remapped the determined T cell subtypes to all the T cells based on clustering analysis (Figures 4A and S5A). In total, we identified 20 T-cell subtypes, including 10 CD4^+^ and 10 CD8^+^ T-cell subtypes using the same criteria previously described (Guo et al., 2018; Zhang et al., 2018; Zhang et al., 2019; Zheng et al., 2017). We found a nice trajectory of both CD4^+^ and CD8^+^ T cells initiated from naïve T cells and further branching into cytotoxic T cells and exhausted T cells (Figures 4A, S4E, and S4H). Immune checkpoint molecules were mainly expressed in regulatory T cells (CD4-c10-FOXP3), pre-exhaustion T cells (CD8-c6-CXCL13), and exhaustion T cells (CD8-c5-PDCD1) as shown in Figure 4B.

**Figure 4.**
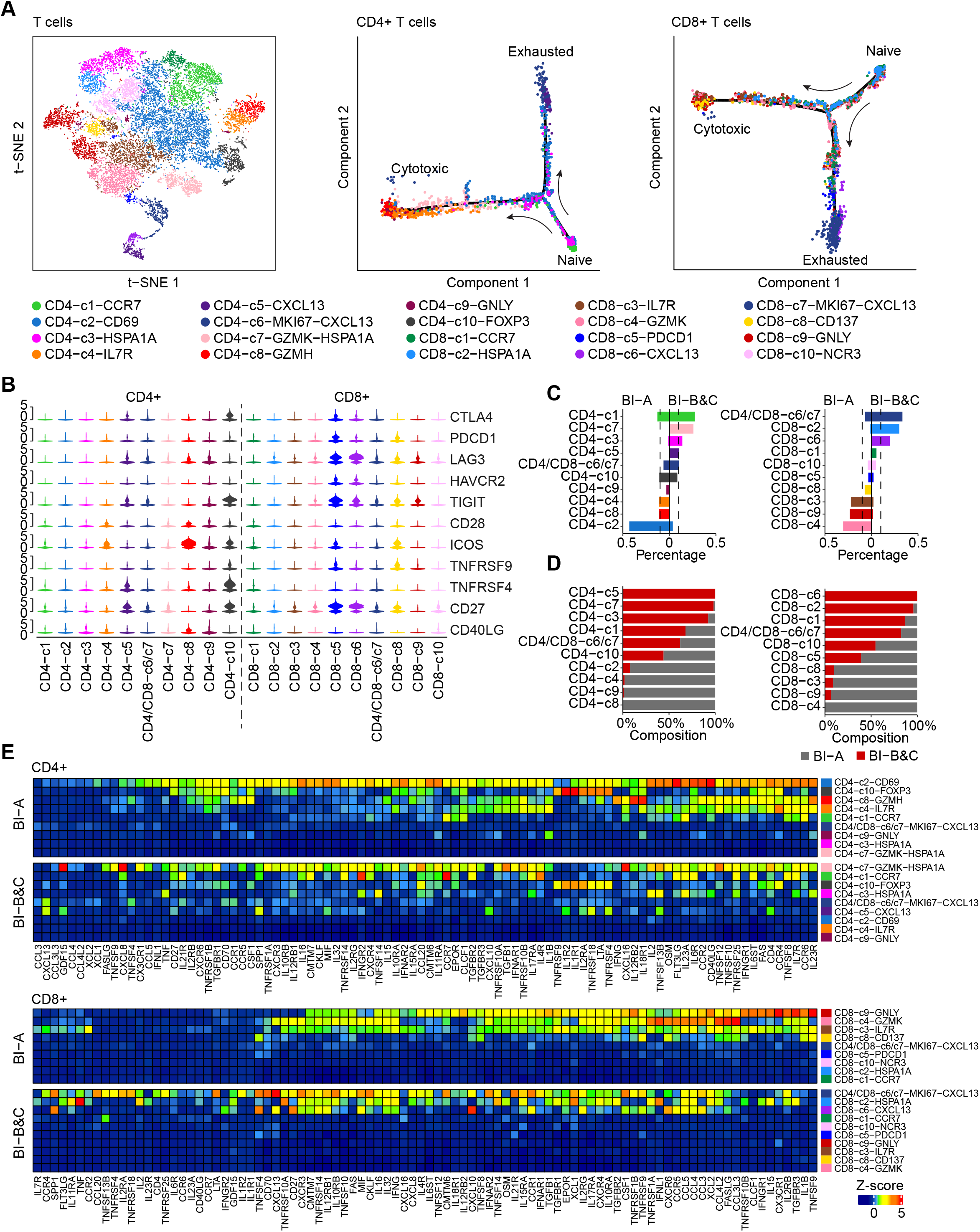
Landscape of T cells in liver cancer. (A) t-SNE plot of all the 19,587 T cells (left), as well as single cell trajectory of CD4^+^ (middle) and CD8^+^ (right) T cells. T-cell subtypes are indicated by colors. (B) Violin plot of immune checkpoint molecules in all T-cell subtypes. The width of a violin plot represents the density of gene expression values. (C) The percentage of each CD4^+^ (left) and CD8^+^ (right) T-cell subtype in BI-A and BI-B&C. Dashed lines indicate 10%. (D) Composition of each CD4^+^ (left) and CD8^+^ (right) T-cell subtype from BI-A and BI-B&C. (E) Cytokines and chemokines produced by each CD4^+^ (top) and CD8^+^ (bottom) T-cell subtype from BI-A and BI-B&C. Gene expression was weighted by the percentage of each T-cell subtype, followed by z-score transformation.

To determine the impact of tumor clonality on T-cell activities, we compared the proportion of T-cell subtypes between BI-A and BI-B&C. We found a polarization of both CD4^+^ and CD8^+^ T-cell subtypes among the two branches (Figures 4C and 4D). Large proportions of memory T cells of CD4-c2-CD69, CD8-c4-GZMK, and CD8-c3-IL7R, as well as a group of cytotoxic T cells of CD8-c9-GNLY were observed in BI-A. In contrast, proliferative pre-exhaustion T cells of CD4/CD8-c6/c7-MKI67-CXCL13 and pre-exhaustion T cells of CD8-c6-CXCL13 were enriched in BI-B&C. In general, the levels of cytokines and chemokines were much higher in CD4^+^ and CD8^+^ T-cells from BI-A than that of BI-B&C (Figure 4E). Among the identified T-cell subsets, cytotoxic T cells of CD8-c9-GNLY were a major source of cytokines and chemokines in BI-A while proliferative pre-exhaustion T cells of CD4/CD8-c6/c7-MKI67-CXCL13 mainly produced those cellular factors in BI-B&C. We found similar results in branching analysis of granzymes and perforin, which may reflect the cytotoxicity of T cells (Figure S5B). We also conducted gene enrichment analysis of CD4^+^ and CD8^+^ T cells from different branches (Figures S5C and S5D). Both CD4^+^ and CD8^+^ T cells from BI-A were enriched in immune response related pathways, which were not found in T cells from BI-B&C. To avoid confounding factors of histologic subtypes of liver cancer, we analyzed T cells from HCC cases separately and similar results were obtained (data not shown). Taken together, these results support a polarization of T cells landscape between BI-A and BI-B&C.

We used a method of CellPhoneDB (Vento-Tormo et al., 2018) to analyze cell communications by searching for ligand-receptor interactions between malignant cells (sources of ligands) and T cells (sources of receptors) to examine potential functional interactions. We observed much stronger interactions between malignant cells and T cells in BI-B&C than that of BI-A for both CD4^+^ and CD8^+^ T cells (Figure 5), suggesting that tumor cells from BI-B&C have an elevated activity of ligand-receptor interactions. Strikingly, SPP1-CD44 was a top interaction pair between malignant cells and T cells, further supporting the key role of SPP1 in the tumor ecosystem.

**Figure 5.**
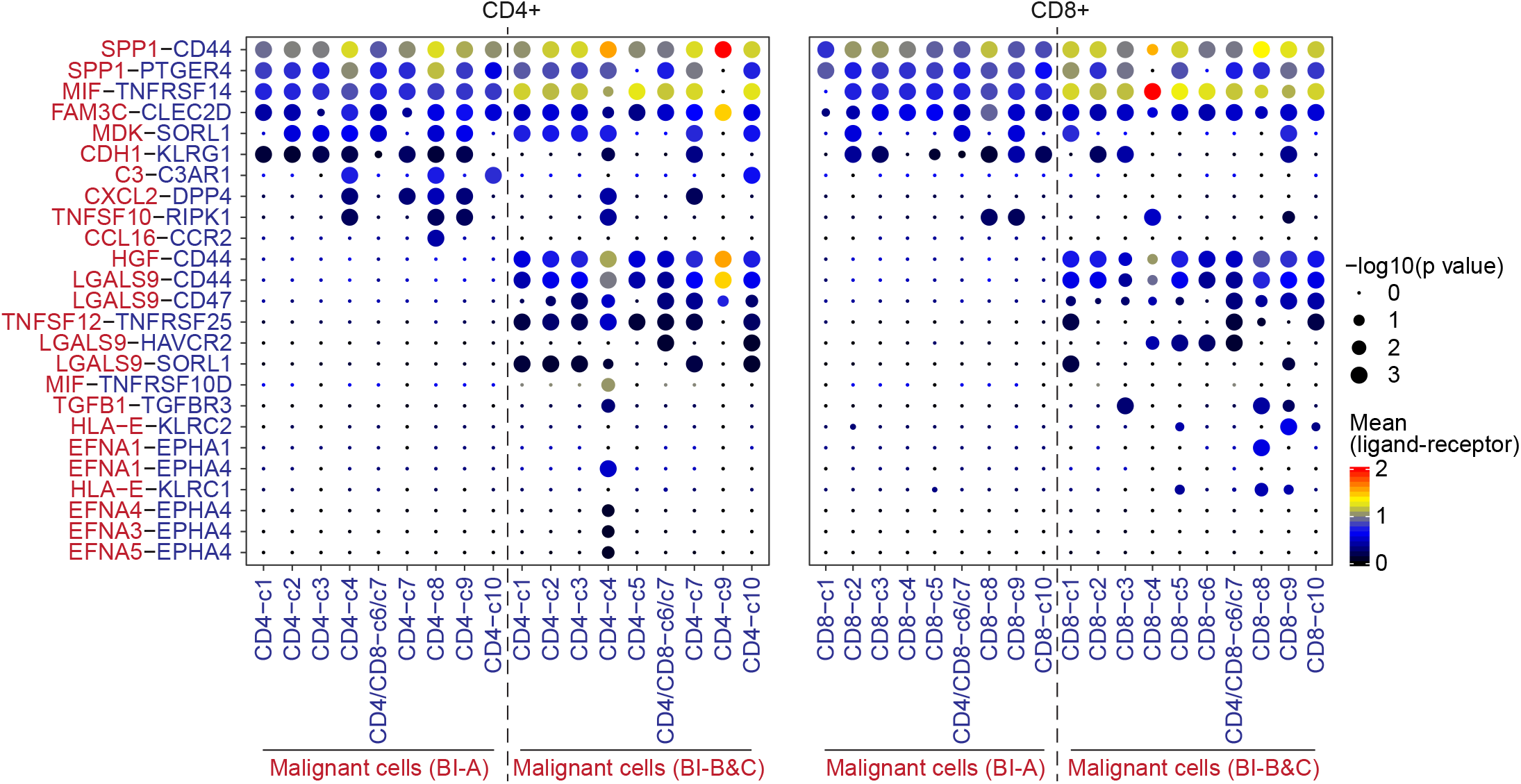
Communications of malignant cells and T cells. Ligand-receptor interactions of malignant cells and CD4^+^ T cells (left) as well as malignant cells and CD8^+^ T cells (right). Dashed lines separate interactions in tumor ecosystems from BI-A and BI-B&C. Row stands for ligand and receptor pairs. red, ligand from malignant cells; blue, receptor from T cells. Column represents pair of malignant cells (red) and T-cell subtype (blue).

### Tumor evolution in response to treatment

Tumor evolutionary trajectories may change over time in response to treatment. Among 37 patients, we obtained at least two core biopsies from different time points for 7 patients (Table S1). Among them, 4 cases with biopsies having at least 15 malignant cells in each biopsy before (baseline) and after treatment were used for evolutionary trajectory analysis by RNA velocity method (Figures 6A and 6B). For two cases of C26 and H68, we found remarkably different lineage of malignant cells before and after treatment, with a weak connection of the clones among two biopsies. In contrast. the clones from before and after treatment in H34 and H73 were mixed, indicating a relatively stable population of malignant cells for these patients during treatment (Figures 6A and 6B). Consistent with the RNA velocity analysis, hierarchical clustering analysis revealed that the clones from baseline and treatment clustered separately for C26 and H68, and were more homogenous in H34 and H73 (Figure 6C). To study genomic similarity of the clones within each case, we calculated pairwise correlation of CNVs between malignant cells before and after treatment (intra-patient) as well as among tumors from different patients (inter-patient). We also performed a randomization of CNV segments to establish a background correlation value distribution as reference. We found a much higher genomic similarity in paired biopsies compared to inter-patient biopsies (Figure 6D), supporting the hypothesis that paired tumors are more likely from the same cell of origin. Interestingly, we found relatively higher degrees of genomic similarity in paired biopsies of C26, H34 and H75, than that of H68 (Figure 6D), suggesting that paired tumor biopsies of H68 are much more genomically distinctive. It is plausible that the follow-up biopsy of H68 may be a *de novo* tumor rather than disseminated tumor cells from its baseline, as its pairwise correlation values are much closer to that of intertumor-derived values compared to C26, H34 and H73 (Figure 6D). Strikingly, SPP1 was significantly elevated in malignant cells of C26 and H68 after treatment, while no significant changes of SPP1 expression was observed in H34 and H73 with similar malignant cell populations before and after treatment (Figure 6E). These results once again highlight the role of SPP1 in tumor evolution.

**Figure 6.**
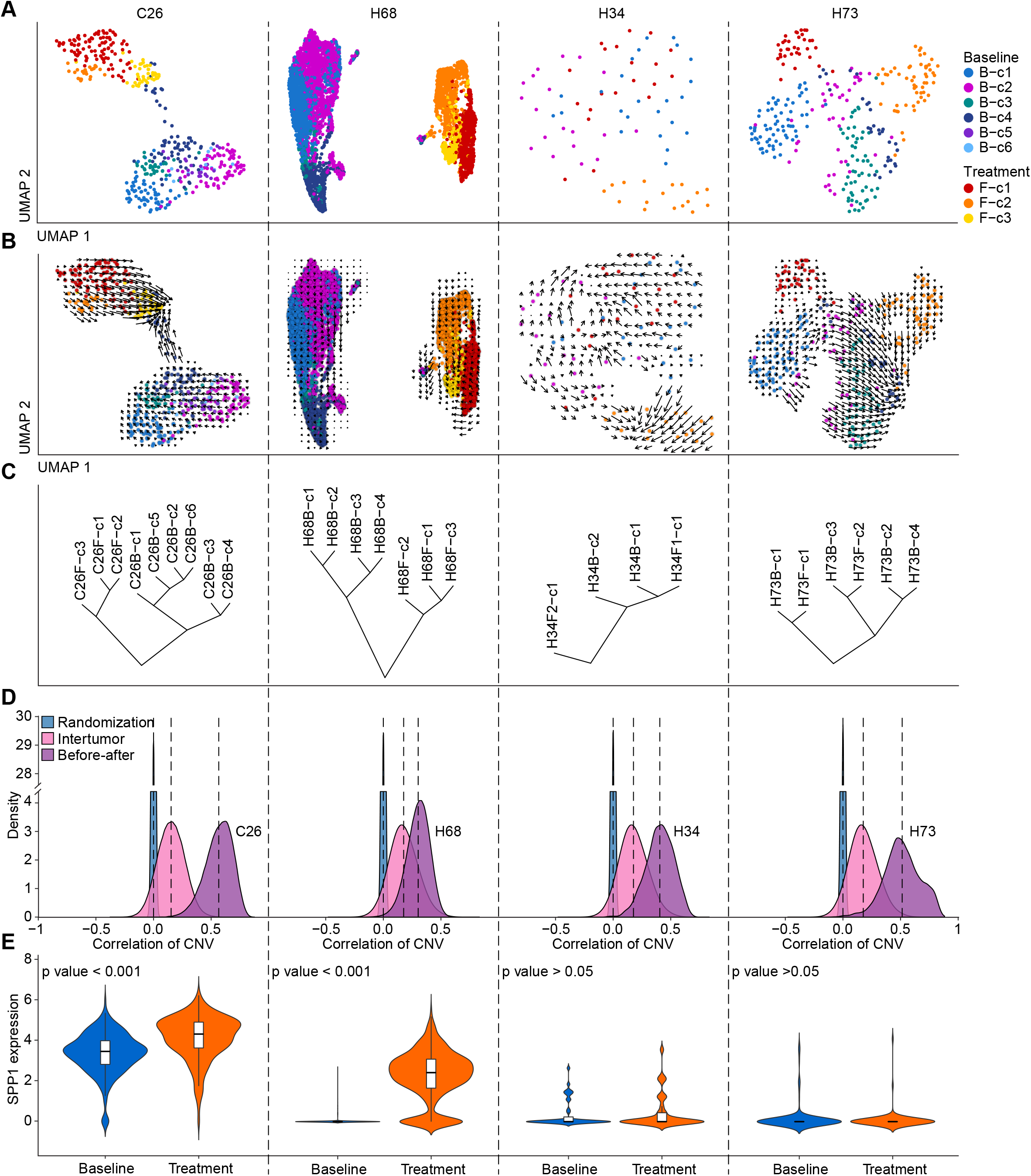
Analysis of liver tumor collected at different time points. (A and B) Tumor cell clonality (A) and RNA velocity-based cell lineage (B) of malignant cells from 4 cases with > 15 malignant cells obtained from each biopsy at baseline or after treatment. Clones are indicated by colors. (C) Hierarchical clustering of the clones from baseline and after-treatment biopsies within each case. (D) Pairwise correlation of CNVs between malignant cells at baseline and after treatment (light purple), between tumors (pink) as well as the correlation of malignant cells with a randomization of CNV segments (blue). (E) SPP1 expression in malignant cells from baseline and treatment. The width of a violin plot represents the density of gene expression values. Box spans the first quartile to the third quartile of the values while segment inside a box indicates the median value.

We also compared the landscape of T cells in biopsies collected between the first biopsy (baseline) and the second biopsy (after treatment) (Figure S5E). There was a smaller difference of CD4^+^ T-cell subtypes among the first biopsy but a much bigger difference among the second biopsy between BI-A and BI-B&C, with a large proportion of memory T cells of CD4-c2-CD69 in BI-A while CD4-c3-HSPA1A enriched in BI-B&C. For CD8^+^ T cells, cytotoxic T cells of CD8-c9-GNLY and proliferative pre-exhaustion T cells of CD4/CD8-c6/c7-MKI67-CXCL13 were enriched in the first biopsy from BI-A and BI-B&C, whereas effector memory T cells of CD8-c4-GZMK were largely in BI-A while CD8-c2-HSPA1A was mainly in BI-B&C from the second biopsy. Consistent with the differential expression patterns of cytokines, chemokines, granzymes and perforin in T cells between BI-A and BI-B&C described above, we also observed similar differential expression patterns between the first biopsy (baseline) and the second biopsy (after treatment) linked to BI-A and BI-B&C (Figure S6). However, it is unclear the functional significance of the T-cell subtypes that express HSPA1A as this subtype has not been described previously (Guo et al., 2018; Zhang et al., 2018; Zhang et al., 2019; Zheng et al., 2017). Further studies are needed to determine the functional roles of CD4-c3-HSPA1A and CD8-c2-HSPA1A T cells in BI-B&C tumors.

## DISCUSSION

Biodiversity is a common trait in solid-organ malignancy and is thought to be evolutionarily selected for its drive to increase tumor cell fitness and survival, possibly via a common molecular mechanism(s). As such, faithfully defining tumor clonality and its evolutionary trajectory is paramount in mechanistic understanding of collective behavior of tumor cells within a solid-organ malignancy and defining its potential drivers. While genome sequencing of bulk tumor has provided compelling evidence for the presence of ITH (Gerlinger et al., 2012), this approach is limited due to its lack of sufficient resolution to define tumor cell clonality with subclones that are functionally distinct and subsequently their distinct functionality-based evolutionary trajectories. The current methods to define driver mutations are also limited for their true coverage since cancer genome is complex, which includes missense mutations, somatic copy number alterations, structural variants and epigenetic alterations. This complexity fuels the ongoing debate about the definitions of tumor clonality and its drivers in the context of tumor heterogeneity (Alizadeh et al., 2015). Improved methodologies are being developed to analyze bulk tumor with the hope to better evaluate tumor clonality (Salcedo et al., 2020). However, given the lack of optimal resolution, it is conceivable that understanding tumor evolution should be sought out at single-cell resolution in a collective effort such as through the NCI Cancer Moonshot Initiative (Rozenblatt-Rosen et al., 2020).

So how do we better define tumor clonality at single-cell resolution, given the current primitive state of single cell sequencing technologies at the genomic level as an error rate for mutational calls is extremely high? We sought to determine liver tumor clonality and its evolution using single cell transcriptomics, an approach that has matured in recent years. Our rationale to use transcriptome to define tumor subclones is as follows. First, while genetic and epigenetic alterations drive tumorigenesis, the vast majority of these alterations may not be linked to phenotypic consequence (Alizadeh et al., 2015). Thus, tracing genetic information with a low degree of confidence in their functional impact could be misleading for monitoring tumor evolution. Second, single cell transcriptomic analysis has been used successfully to define distinct cell types with defined functionality-based clones and phenotypic features in various organs such as brain and liver (Aizarani et al., 2019; Tasic et al., 2018; Zhong et al., 2018). Third, each tumor is comprised of cell populations with different transcriptional programs. As tumor subclones evolve through phenotypic selection to adapt to various environments, functional diversity may be the consequence of genetic heterogeneity. Thus, tracing subclones based on their transcriptional program may represent a closer cellular state in regards to their phenotypic features, a strategy that has been increasingly recognized as tumor functional heterogeneity (Gonzalez-Silva et al., 2020). Using this strategy, we have determined tumor cell clonality and its hierarchical relationship in HCC and iCCA. We found that patients whose tumors have a higher clone number have a much shorter survival compared to those with a lower clone number. Phylogenetic analysis of subclones reveals strong evidence of functionality-based tumor branching evolution in HCC and iCCA, consistent with the Darwinian tumor evolution model of Peter Nowell (Nowell, 1976). CNV analysis revealed similarities in clonality between transcriptome-based subclones and CNV-based subclones, supporting the hypothesis that a functional diversity is the consequence of genetic heterogeneity and thus better represents functionality-based evolutionary trajectories. Moreover, we validated our findings using independent and publicly available tumor bulk data from 488 HCC patients and 277 iCCA patients. Interestingly, iCCA generally has a much higher clonality than HCC among patients enrolled into our interventional studies and it is also more refractory to treatment than HCC. It is plausible that tumor cell evolution may drive an increase in biodiversity thus rendering the treatment ineffective.

Identification of key players driving tumor cell evolution is of significant importance as it can facilitate the development of better approaches for cancer intervention. Using two independent strategies, including searching conserved genes among different branches and ligand-receptor interaction networks that may be responsible for tumor cell evolution, we identified SPP1 as the top candidate player. Consistently, we found that the SPP1 expression patterns follow hierarchical relationships of tumor cell branching evolution with a much higher expression level in BI-B/C clones than BI-A clones. Moreover, SPP1 is significantly elevated in tumors after treatment in C26 and H68 but not H34 and H73. Both H34 and H73 are considered responders in our intervention studies with a good prognosis and they both belong to BI-A tumors. While H68 is also included as BI-A tumor, SPP1 is elevated in the second biopsy of this patient. Pairwise CNV analysis of the first and the second biopsies of this patient revealed a much lower genomic similarity compared to before and after treatment biopsies from other patients. Consistently, RNA velocity analysis also showed a lack of connectivity between before and after biopsies of H68, raising the possibility that the second biopsy of H68 may be a *de novo* tumor. It is well known that SPP1 encodes osteopontin (OPN), a phosphorylated glycoprotein expressing in various tissues and cell types linked to human diseases (Icer and Gezmen-Karadag, 2018; Rittling and Singh, 2015). The role of OPN in tumor progression including HCC metastasis and prognosis is well documented (Rittling and Chambers, 2004; Ye et al., 2003). Its ability to induce tumor cell invasion and metastasis in HCC is likely to be linked to its ability to drive tumor cell evolution that can adapt tumor microenvironment for survival fitness. Furthermore, OPN may regulate different types of immune cells by inducing the reprogramming of these cells towards innate state against pathogens, thereby reducing an effective host immunity. OPN exists as several isoforms that exert different functions by interacting differentially with its receptors CD44 and integrins (Briones-Orta et al., 2017; Kanayama et al., 2017; Takafuji et al., 2007). Collectively, our results indicate that SPP1 may functionally drive tumor cell evolution in HCC and iCCA, possibly by reprogramming tumor immunity by damping the host immune response to tumor cells. However, the functional link between SPP1 and tumor cell evolution remains to be tested in an experimental model system. Given the key roles of OPN in tumor progression, rational design such as the development of neutralizing antibodies specific to certain isoforms of OPN is warranted.

As cancers change over time through a process of tumor cell evolution, the ability to characterize tumors during their progression and its response to treatments is the key to better defining tumor biology and developing more effective therapy (Maley et al., 2017). Using single cell transcriptome analysis of biopsies from HCC and iCCA patients before and after treatment, we have successfully obtained the tumor cell landscape for 37 patients and constructed tumor cell evolutionary trajectories. We identified SPP1 as a candidate driver for tumor cell evolution and determined potential mechanisms for HCC and iCCA patients who may not respond to treatments such as immune checkpoint blockade. Although tumor biopsy is not routinely used in HCC diagnosis, classification based upon biopsy may inform a diagnostic stratification for therapy or prognosis, a strategy currently being implemented into the NCI Liver Cancer Moonshot program. These extra steps may help to explain why some patients respond to immunotherapy while others do not. Discovery and validation of non-invasive surrogate imaging biomarker correlates of clonality density could pinpoint evolutionary trends when they occur, instead of awaiting for the poor outcomes themselves. Deciphering tumor and cellular heterogeneities within liver cancers could also guide which index lesions or clonal volumes to locally ablate, versus others immunomodulate with image guided local therapies. Development of such a classification system holds promise for delivering better precision medicine with the benefit to improve life of patients with liver cancer.

## METHODS

### Human sample collection

Fresh liver tumor biopsies were collected with preoperative informed consent from patients participating at NIH Clinical Center for interventional studies, following approval by the ethics committee of the National Institutes of Health. A total of 37 liver cancer patients (HCC, 25; iCCA, 12) were involved in this study, with 46 tumor biopsies collected before or after treatment. Among the tumor biopsies, 19 samples were collected and analyzed in our previous study (Ma et al., 2019). The ages of the patients ranged from 35 to 81, with a median age of 65. Fourteen patients were tested positive for HCV, four patients positive for HBV, two with fatty liver and one positive for both HBV and HDV. A majority of the patients were treated with immune checkpoint inhibitors of durvalumab, tremelimumab, pembrolizumab to target PD-L1, CTLA-4 as well as PD-1. The detailed clinical data of the patients was summarized in Table S1.

### Tissue dissociation

Core needle biopsies or resected tumor tissues were collected in MACS Tissue Storage Solution (Miltenyi Biotech, Cat# 130-100-008) and immediately dissociated on ice to minimize ischemic time. Tissues were cut into small pieces of 2-4 mm with a sterile scalpel after removing fat, fibrous and necrotic areas. Then they were transferred to a gentleMACS C Tube (Miltenyi Biotech, Cat#130-093-237) with 5ml enzymatic digestion mix (Tumor dissociation Kit from Miltenyi Biotech, Cat# 130-095-929) for homogenization by using a gentleMACS Dissociator (Miltenyi Biotech). Homogenized tissues were enzymatically digested to single cells at 37 °C for 30 min with 300 rpm shaking. Undigested tissues were removed using a 70-um cell strainer (Miltenyi # 130-095-823). Single cells were finally collected by centrifuge at 300xg and 4 °C for 5 min for further single-cell capture or long-term storage in liquid nitrogen.

### Library preparation and scRNA-seq

Single-cell cDNA library was constructed following the 10x Genomics Single Cell 3’ Reagent Kit v2 or v3 user guide. Single-cell suspension was prepared and adjusted to 1,200 cells/μl for Gel Bead-In-EMulsion (GEM) generation and cell barcoding. In each GEM, a single cell, a single gel bead, as well as reverse transcription (RT) reagents were involved for RT of mRNA. Post GEM-RT cleanup and cDNA amplification were then performed, followed by single-cell library construction. Illumina HiSeq 4000/NovaSeq 6000 system was used for cDNA library sequencing, with parameters of 26/28 cycles for barcodes, 8 cycles for sample indices and 98 cycles for cDNA reads. The sequencing depth was targeted as 50,000 raw reads per cell. Base calling was carried out with Real-Time Analysis software on Illumina sequencing systems. Demultiplexing was then performed by using bcl2fastq, with one mismatch allowed in the barcodes. Cell Ranger was further applied for alignment, tagging, gene and transcript counting. All samples had excellent sequencing yield, with raw reads ranging from 105–465 million per sample and > 95% Q30 bases in barcode. The scRNA-seq data were submitted to the Gene Expression Omnibus (accession number GSE151530).

### scRNA-seq data processing

We first applied Cell Ranger (version 3.1.0) to perform sequencing depth normalization and chemistry batch correction of all the tumor samples. Then Seurat (version 3.1.2) (Butler et al., 2018) was used for initial data analysis. We removed the genes that were expressed in < 0.1% of all the single cells, and further eliminated the cells with < 500 detected genes. After the initial quality control, a total of 56,721 cells were retained. We normalized the raw counts in each cell by the total number of counts and multiplied by a scale factor of 10,000, followed by a log transformation. The top 2,000 highly variable genes were selected by using variance-stabilizing transformation method (Stuart et al., 2019). Those genes represented the heterogeneous features of the single cells for downstream dimensional reduction and data visualization. We further applied a linear transformation to the selected genes, as a pre-processing step prior to principal component analysis (PCA). According to eigenvalues from PCA, we used the first 10 principal components (PCs) to perform t-distributed stochastic neighbor embedding (t-SNE) analysis of all the single cells.

### Identification of malignant cells and non-malignant cells

We separated malignant cells and non-malignant cells by estimating CNV from single-cell transcriptome, as previously described (Ma et al., 2019). A slide window of 100 genes was applied for calculating the average relative expression of genes to eliminate gene-specific patterns and to reflect the CNV. We defined CNV score (the sum of the squared CNV profiles of a single cell) and CNV correlation score (the correlation of CNV of a single-cell and the average CNV profiles of the top 2% single cells in CNV score within a tumor) to discriminate malignant cells from non-malignant cells. To further confirm the successful separation of malignant cells and non-malignant cell, we calculated epithelial score and liver marker score, which were defined as the average expression of 14 epithelial marker genes (Puram et al., 2017) and 24 liver marker genes (Kim et al., 2017). We also performed doublet removal of single cells following a method described in a recent study (Tasic et al., 2018).

### Clonality determination of malignant cells

We developed a consensus clustering method to determine clones of malignant cells within a tumor. We used a sampling strategy to ensure the fairness of clonal detection from the tumor samples with imbalanced number of malignant cells. Machine learning algorithms and statistical methods were further applied to refine the detected clones. A schematic overview of the workflow was provided in Figure S2A. Below, we listed the detailed steps of clonality determination:

1. Sampling and initial clustering. For a tumor sample with > 200 malignant cells, we performed random sampling of malignant cells (n = 200) within the malignant-cell population. Then we conducted clustering of the sampled malignant cells by using SC3 (Kiselev et al., 2017), a clustering method designed for scRNA-seq data. We used the estimated optimal clustering provided by SC3 as the initial clusters in our analysis.
2. Cluster refinement. We performed PCA of the sampled malignant cells based on the top 1000 highly variable genes. Then for each pair of clusters determined in step 1, we trained a support vector machine (SVM) model by using the first 10 PCs. Ten-fold cross-validation was carried out to calculate the accuracy of the cluster prediction. Then we performed 100 permutations by shuffling the labels (clusters) of the single cells. Based on the accuracy of the original cluster prediction as well as the accuracies from the permutation test, a permutation p value for a pair of selected clusters can be derived. After generating the permutation p values of all possible pairs of clusters, we ranked the p values from the largest to the smallest. If the largest permutation p value was greater than 0.05, the corresponding pair of clusters would be merged as one cluster. With the newly generated clusters, we repeated the process of cross-validation and permutation test for each pair of clusters until the largest permutation p value < 0.05.
3. Cluster determination of all the malignant cells within a tumor. With the refined clusters of the sampled cells, we trained an SVM model based on the first 10 PCs. For the rest of the malignant cells within a tumor, we first projected those single cells to the PCA space of the sampled cells. Based on the projected PCs, we predicted the cluster information of those single cells by using the well-trained SVM model. Then, the cluster information of all the malignant cells within a tumor would be determined.

For a tumor sample with > 200 malignant cells, we repeated steps 1‒3 for 100 times. We then determined the number of clusters within a tumor based on the statistics of clusters obtained from each iteration. Finally, a consensus matrix was calculated by using the results from the iteration process to determine the final cluster information of all the malignant cells. For a tumor sample with ≤ 200 malignant cells, we directly performed clustering of all the malignant cells using SC3, followed by cluster refinement in step 2 to determine the clusters within a tumor.

Based on the approach described above, the number of clusters derived from tumors with < 200 malignant cells were not comparable those obtained from tumors with > 200 malignant cells. Thus, we inferred the number of clusters for this group of tumor samples. We used the samples with > 200 malignant cells as a reference group, and the samples with < 200 malignant cells as a testing group. For each tumor in the testing group, we calculated the pair-wise correlation of malignant cells between the selected tumor and all the tumors in the reference group. Then the number of clusters in the selected tumor was inferred as that within the most similar tumor—the one with the highest mean correlation score. The inferred number of clusters was only used for performing comparison of the number clusters between tumors.

### Construction of the phylogenetic tree

We used hierarchical clustering method to construct the phylogenetic tree of the clones from all the tumors. The top 200 most variable genes within each tumor were selected, considering the fairness of tumors with imbalanced number of malignant cells. We then removed mitochondrial genes and ribosomal genes (Ramachandran et al., 2019). A total of 2,555 unique genes were derived and assembled as a gene list for hierarchical clustering. We used the median expression of these genes in each tumor clone as its centroid. A distance matrix of the clones was further calculated, based on which hierarchical clustering was carried out with complete linkage. We used bootstrap method (1000 bootstrap replications) in R package pvclust (version 2.2.0) to calculate p values and measure the confidence of the hierarchical relationship of the clones.

### Single-cell trajectory analysis

We used Monocle package (version 2.14.0) (Trapnell et al., 2014) to learn single-cell trajectory of malignant cells to independently determine the clonal relationship within a tumor. The genes expressed in at least 10% of all the malignant cells within a tumor were applied for PCA and t-SNE analysis, where clusters were identified by density peak clustering method. With the differential expression genes among clusters, we employed reverse graph embedding method to learn the tree-like manifold and to construct single-cell trajectory (Qiu et al., 2017). We also used this method to learn the trajectories of CD4^+^ T cells as well as CD8^+^ T cells. We excluded CD4-c10-FOXP3 (regulatory T cells) and CD8-c10-NCR3 (largely composed of mucosal associated invariant T cells) from the trajectory analysis due to their distinct characteristics, following a strategy used in a recent study (Guo et al., 2018).

### RNA velocity-based cell lineage tracing

We traced cell lineage of malignant cells within each tumor by using RNA velocity as another independent way to study tumor clonal relationship. A python tool of velocyto.py was applied to annotate the spliced and unspliced reads from the bam files of scRNA-seq data to generate the loom files. We estimated RNA velocity of malignant cells by using *RunVelocity* function in R package SeuratWrappers (version 0.1.0, wrapped velocyto.R), and further visualized the velocity on a UMAP by calculating a correlation-based transition probability matrix within a k-nearest neighbor graph.

### Conserved genes with tumor branching evolution

We searched for conserved genes of different branches to find possible drivers of tumor branching evolution. We started from the top 200 highly variable genes of malignant cells within a tumor, and selected genes that were presented in ≥ 50% clones, with gene expression in ≥ 20% cells of each clone. A group of conserved genes among clones within each tumor can be obtained during this process. We further selected the genes that were presented in ≥ 50% samples of a tumor branch to generate the conserved genes at branch level. We also performed *t*-test of the genes in all the clones of the tumors from different branches, to ensure the selected genes were branch specific. Considering a small number of cases in BI-B, we merged samples in BI-B and BI-C to avoid false positive hits in searching for conserved genes.

### Gene enrichment analysis

Gene enrichment analysis was carried out on hallmark gene sets by using GSEA (version 4.0.3) (Subramanian et al., 2005). For malignant cells, we performed differential gene expression analysis among the clones within each tumor. Then the derived differential expression genes were used as input of GSEA. Thus, only the tumors with at least two clones detected were involved in this part of analysis. For T cells, we applied the genes that were expressed in at least 10% of all the CD4^+^ or CD8^+^ T cells to find the enriched pathways in T cells from BI-A or BI-B&C. We performed 1000 permutations in the analysis to generate the statistics. Pathways with FDR adjusted q value < 0.05 were selected for plot generation.

### Survival risk prediction using bulk transcriptomic data

We applied BRB-ArrayTools (version 4.6.0) (Simon et al., 2007) to perform overall survival analysis of patients in ICGC cohort (CCA, n = 115), Japan cohort (CCA, n = 162), TCGA cohort (HCC, n = 249), and LCI cohort (HCC, n = 239). We carried out differential gene expression analysis of malignant cells from all cases or only HCC cases in BI-A and BI-B&C, in order to find surrogate gene signatures of the two branches. A cox proportional hazards model was built upon the first 10 PCs of the surrogate gene signature from BI-A or BI-B&C for each cohort. A total of 100 permutations were performed for log-rank test of low-risk and high-risk groups of patients, with log-rank p value and permutation p value provided to indicate the statistical significance.

### Identification of T cell subtypes

To ensure a pure population of T cells, we calculated the proportion of T cells expressing CD3D, CD3E, or CD3G in each cluster generated from a graph-based clustering algorithm with default parameter settings (Waltman and Van Eck, 2013). The clusters with ≤ 80% T cells were removed from further analysis (Figure S4B). To confidently identify T cell subtypes, we first extracted CD4^+^ (expressing CD4, n = 2,275) T cells and CD8^+^ (expressing CD8A or CD8B, n = 6,686) T cells, respectively. Then for each group of T cells, we constructed a shared nearest neighbor graph based on the k-nearest neighbors of each cell. Louvain algorithm was further applied to optimize the modularity function to determine clusters. We performed differential gene expression analysis of the derived clusters and named the T cell subtypes according to known marker genes (Figures S4C and S4F). We also evaluated each T cell subtype by using the 4 well-defined naïve T cell markers (CCR7, LEF1, SELL, TCF7), 12 cytotoxic markers (PRF1, IFNG, GNLY, NKG7, GZMB, GZMA, GZMH, KLRK1, KLRB1, KLRD1, CTSW, CST7) as well as T-cell exhaustion marker genes (Figures S4D and S4G) (Guo et al., 2018). Exhaustion marker genes were generated by comparing pre-exhaustion and exhaustion T cells with all other T-cell subtypes, and further filtered by CD4^+^ or CD8^+^ T-cell exhaustion related gene signatures provided in a recent study (Guo et al., 2018). We determined the subtypes of all the T cells (n = 19,587) by remapping the well-defined T-cell subtypes based on clustering analysis (Figure S5A). Specifically, we performed clustering of all the T cells using the aforementioned graph-based algorithm and obtained 39 T-cell clusters by setting the parameter of *resolution* = 3 in order to generate a larger number of clusters. Then we calculated the proportion of the well-defined CD4^+^ and CD8^+^ T-cell subtypes in each T-cell cluster. The T-cell subtype of a specific cluster was determined as the most abundant well-defined T-cell subtype in that cluster. In all the derived clusters, there was one dominant well-defined T-cell subtype, indicating the successful identification and mapping of T-cell subtypes (Figure S5A).

### Communications of malignant cells and T cells

We used CellphoneDB (Vento-Tormo et al., 2018) to study the ligand-receptor interactions of malignant cells and T cells. We carried out 1000 permutations to generate the null distribution of the mean of the ligand expression in one cell type and the receptor expression in another cell type. Any ligand-receptor pairs with p value < 0.05 were considered as significant. To study the reprogramming of T cells by secreted cellular factors from malignant cells, we filtered the interaction pairs by selecting those with ligand produced by malignant cells and receptor provided by T cells, followed by removing the pairs with integrin detected.

### Correlation of CNV profiles

We calculated the Pearson’s correlation of the CNV profiles of malignant cells from biopsies collected at baseline and during treatment of the same patient to indicate the tumor evolution in response to treatment. A density plot of all the correlation coefficients from one case was generated by the *geom_density* function in R package ggplot2 (version 3.2.1). The correlation of CNVs of the malignant cells between HCC tumors and between CCA tumors was calculated as reference for HCC and CCA cases, respectively. We also performed randomization of the CNV segments of two randomly selected malignant cells for correlation calculation. The process was repeated for 100,000 times to generate a density plot of the correlation coefficients to establish a background.

### Histopathology

We obtained the histopathology images from the Laboratory of Pathology at National Cancer Institute. Tumor biopsies were sectioned at 5μm and routinely stained with hematoxylin and eosin for further examination.

### Data availability

Single-cell transcriptomic data are available at the Gene Expression Omnibus (accession number GSE151530).

## ACKNOWLEDGEMENTS

We thank members of the Wang laboratory for critical discussions; the Greten laboratory for managing clinical programs; Sean P. Martin for clinical data collection; Wei Tang for advice on data analysis; Zachary Rae for additional laboratory work; the patients, families and nurses for contribution to this study. This work was supported by grants (ZIA BC 010877, ZIA BC 010876, ZIA BC 010313 and ZIA BC 011870) from the intramural research program of the Center for Cancer Research, National Cancer Institute of the United States.

## AUTHOR CONTRIBUTIONS

X.W.W. developed study concept; T.F.G. directed clinical study; L.M. and X.W.W. directed experimental design and interpreted data; L.M. performed computational analysis; L.W., C.W.C., S.F., D.D., M.F., J.C., M.O.H., M.K., Y.Z., B.T., J.M.H., J.L.D., D.E.K., B.J.W., T.F.G. conducted experiments and additional data analysis; L.M. and X.W.W. wrote the manuscript with help from L.W. All authors read, edited, and approved the manuscript.

## SUPPLEMENTARY FIGURE LEGENDS

**Figure S1.**
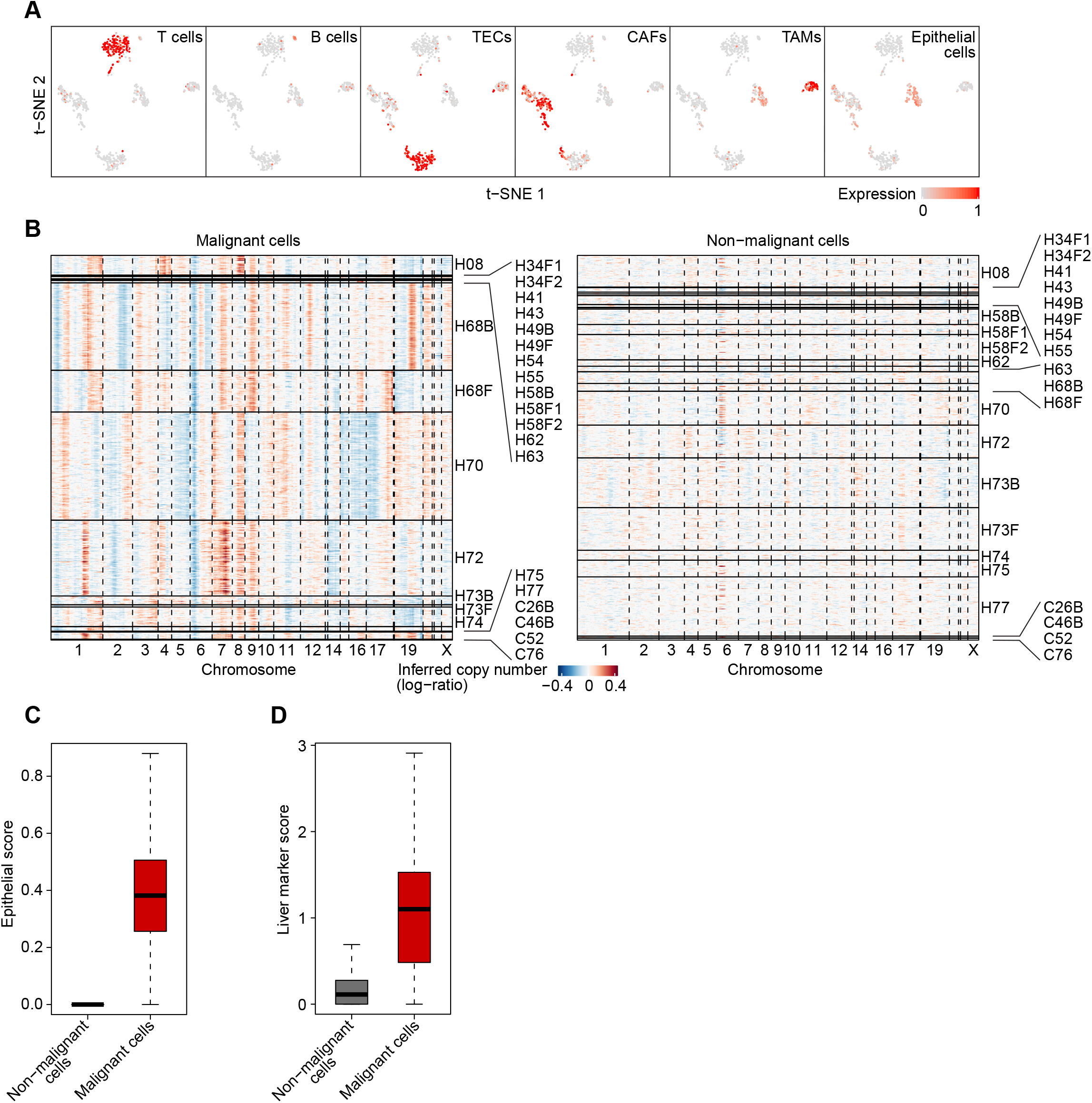
Separation of malignant cells and non-malignant cells. (A) Annotation of single cells from H34. Cells were annotated based on average expression of known cell lineage-specific marker genes of T cells (CD2, CD3E, CD3D, CD3G), B cells (CD79A, SLAMF7, BLNK, FCRL5), TECs (PECAM1, VWF, ENG, CDH5), CAFs (COL1A2, FAP, PDPN, DCN, COL3A1, COL6A1), TAMs (CD14, CD163, CD68, CSF1R) as well as epithelial cells (KRT8, KRT15, KRT16, KRT17, KRT18, KRT19, SFN, KRTCAP3, EPCAM). (B) Large-scale CNVs of malignant cells (left) and non-malignant cells (right) inferred from single-cell transcriptome. Red, amplification; blue, deletion. (C and D) Epithelial score (C) and liver marker score (D) of malignant cells and non-malignant cells. Box spans the first quartile to the third quartile of the values while segment inside a box indicates the median value.

**Figure S2.**
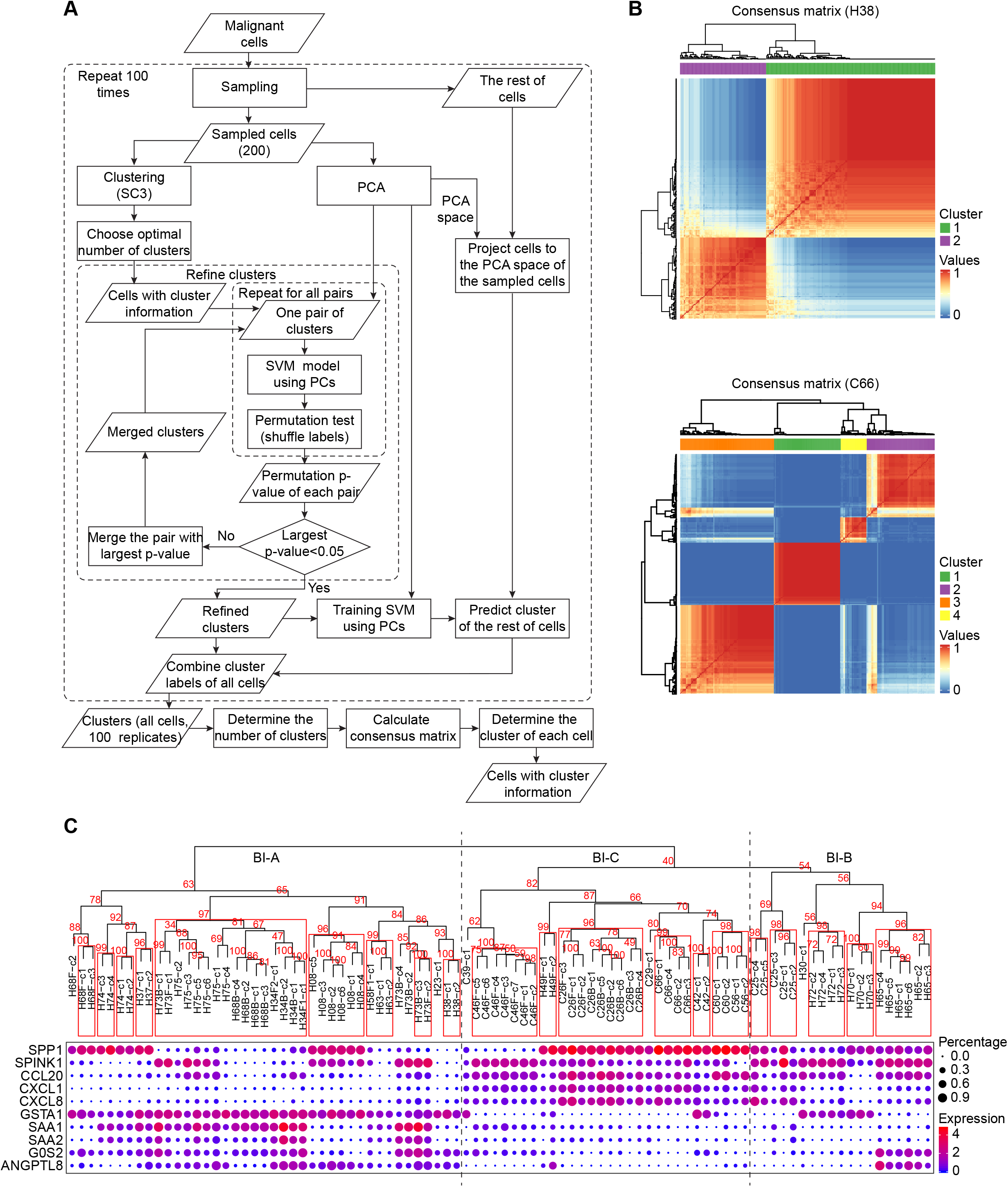
Determination of tumor cell clonality. (A) Schematic overview of tumor cell clonality determination of malignant cells. (B) Consensus matrix of clustering of malignant cells from two representative tumors of H38 and C66 by using the strategy in (A). (C) Top panel shows the bootstrapped confidence of the hierarchical relationship of the clones from all tumors. Bottom panel lists the conserved genes of different branches. Approximately unbiased (AU) p-value (%) computed by multiscale bootstrap resampling was provided, with AU > 95% highlighted by rectangles.

**Figure S3.**
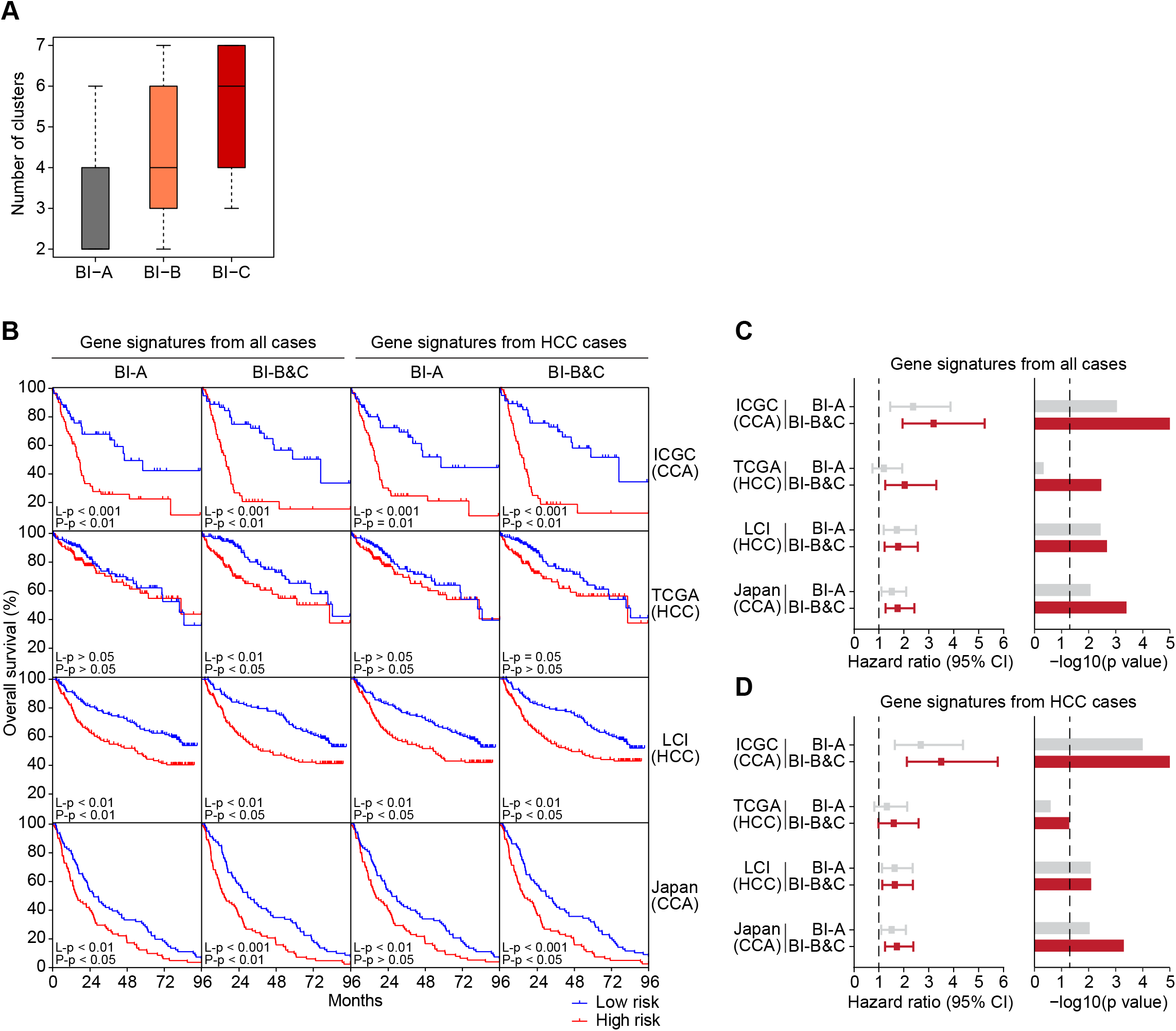
Survival risk prediction based on tumor branching related genes. (A) The number of clones detected from tumors in BI-A, BI-B and BI-C. Box spans the first quartile to the third quartile of the values while segment inside a box indicates the median value. (B) Kaplan-Meier plots of the overall survival of ICGC (CCA), TCGA (HCC), LCI (HCC) and Japan (CCA) cohorts based on tumor branching-specific gene signatures generated by comparing malignant cells from BI-A and BI-B&C. The gene signatures were obtained from all tumors, as well as only HCC tumors in the phylogenetic tree. Low risk and high-risk groups are colored blue and red, respectively. Log-rank p value (L-p) and permutation p value (P-p) of the survival analysis were provided. (C and D) Hazard ratio with 95% confidence interval (CI) and log-rank p value derived from survival risk prediction in (B) using gene signatures from all cases (C) or only HCC cases (D). Dashed lines indicate hazard ratio =1 or log-rank p value = 0.05.

**Figure S4.**
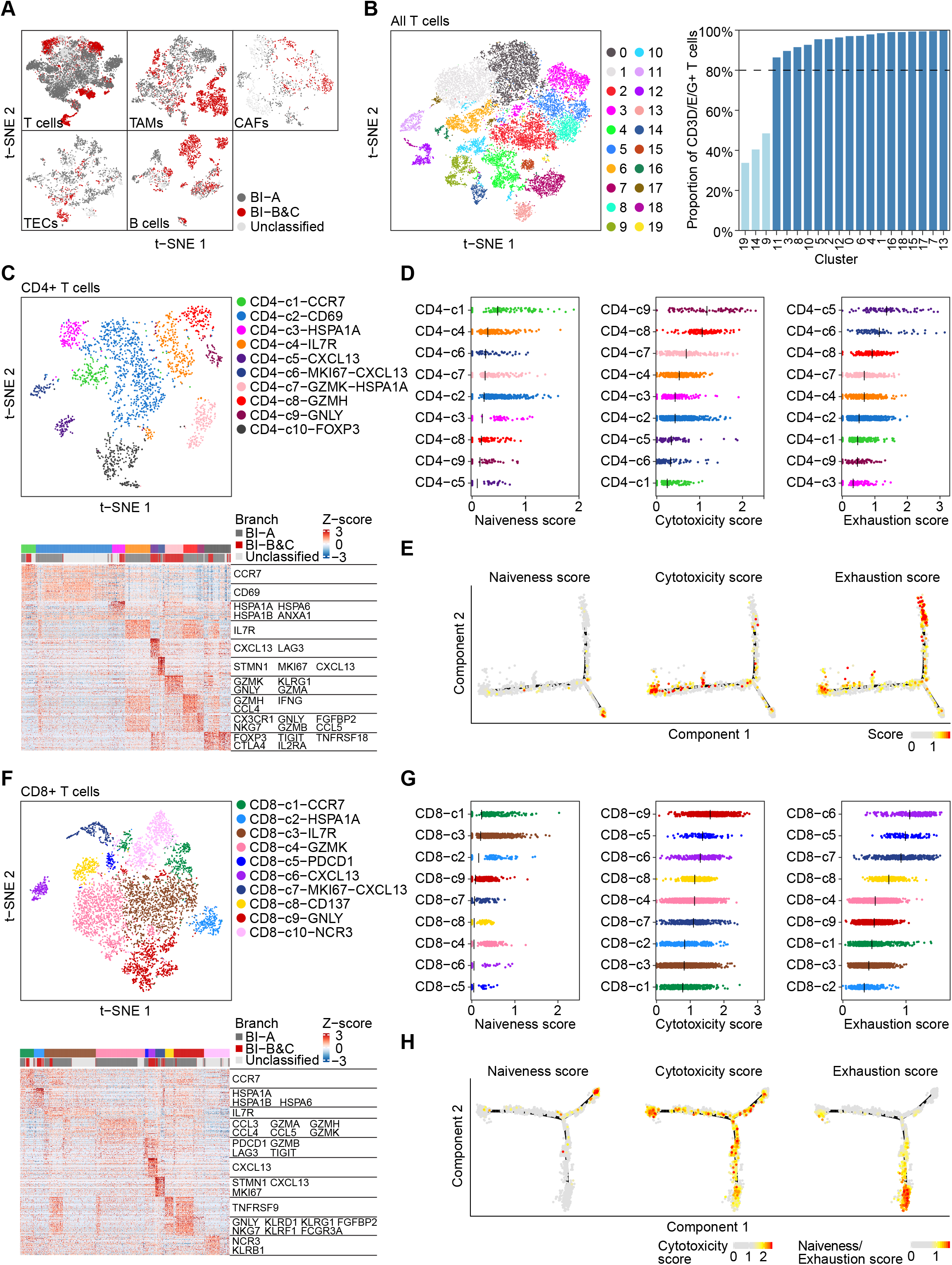
Determination of T-cell subtypes. (A) t-SNE plot of T cells, TAMs, CAFs, TECs, and B cells colored by tumor branching information. The single cells that were derived from samples with ≤ 15 malignant cells (not involved in the tumor phylogenetic tree) were labeled as unclassified. (B) Clustering of all T cells (left), and the proportion of T cells expressing CD3D/E/G in each derived cluster (right). (C) Clustering (top) of CD4^+^ T cells and known marker genes (bottom) of each CD4^+^ T-cell subtype. (D) Naïveness score (left), cytotoxicity score (middle) and exhaustion score (right) of each CD4^+^ T-cell subtype. Mean values are indicated with line segments. (E) Single-cell trajectory of CD4^+^ T cells colored by naïveness score, cytotoxicity score and exhaustion score. (F) Clustering (top) of CD8^+^ T cells and known marker genes (bottom) of each CD8^+^ T-cell subtype. (G) Naïveness score (left), cytotoxicity score (middle) and exhaustion score (right) of each CD8^+^ T-cell subtype. Mean values are indicated with line segments. (H) Single-cell trajectory of CD8^+^ T cells colored by naïveness score, cytotoxicity score and exhaustion score.

**Figure S5.**
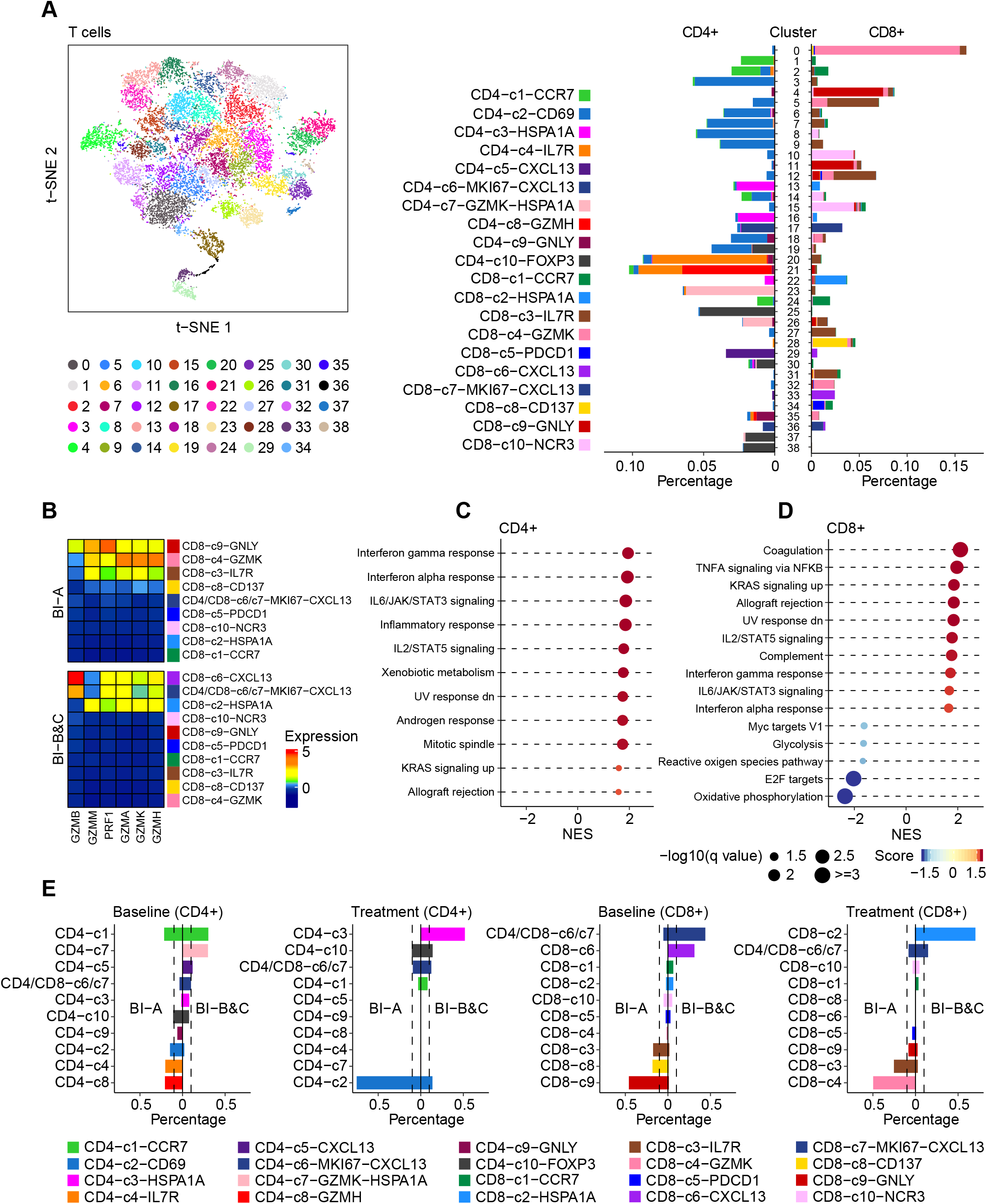
Analysis of T cells. (A) Clustering (left) of T cells and the proportion (right) of T-cell subtypes in each of the T-cell cluster. The percentage of each CD4^+^ T-cell subtype as well each CD8^+^ T-cell subtype were calculated through all the T-cell clusters to determine the T-cell subtype of each cluster. (B) Granzymes and perforin produced by T cells from BI-A and BI-B&C. Gene expression was weighted by the percentage of each T-cell subtype, followed by z-score transformation. (C and D) Gene enrichment analysis of CD4^+^ (C) and CD8^+^ (D) T cells from BI-A and BI-B&C. A positive NES (red) represents the corresponding pathway was enriched in T cells from BI-A, while a NES (blue) stands for a pathway enrichment in BI-B&C. NES, normalized enrichment score. (E) Comparison of the proportion of each T-cell subtype derived from BI-A and BI-B&C at baseline or after treatment. Panels from left to right: CD4^+^ T cells at baseline, CD4^+^ T cells after treatment, CD8^+^ T cells at baseline, and CD8^+^ T cells after treatment.

**Figure S6.**
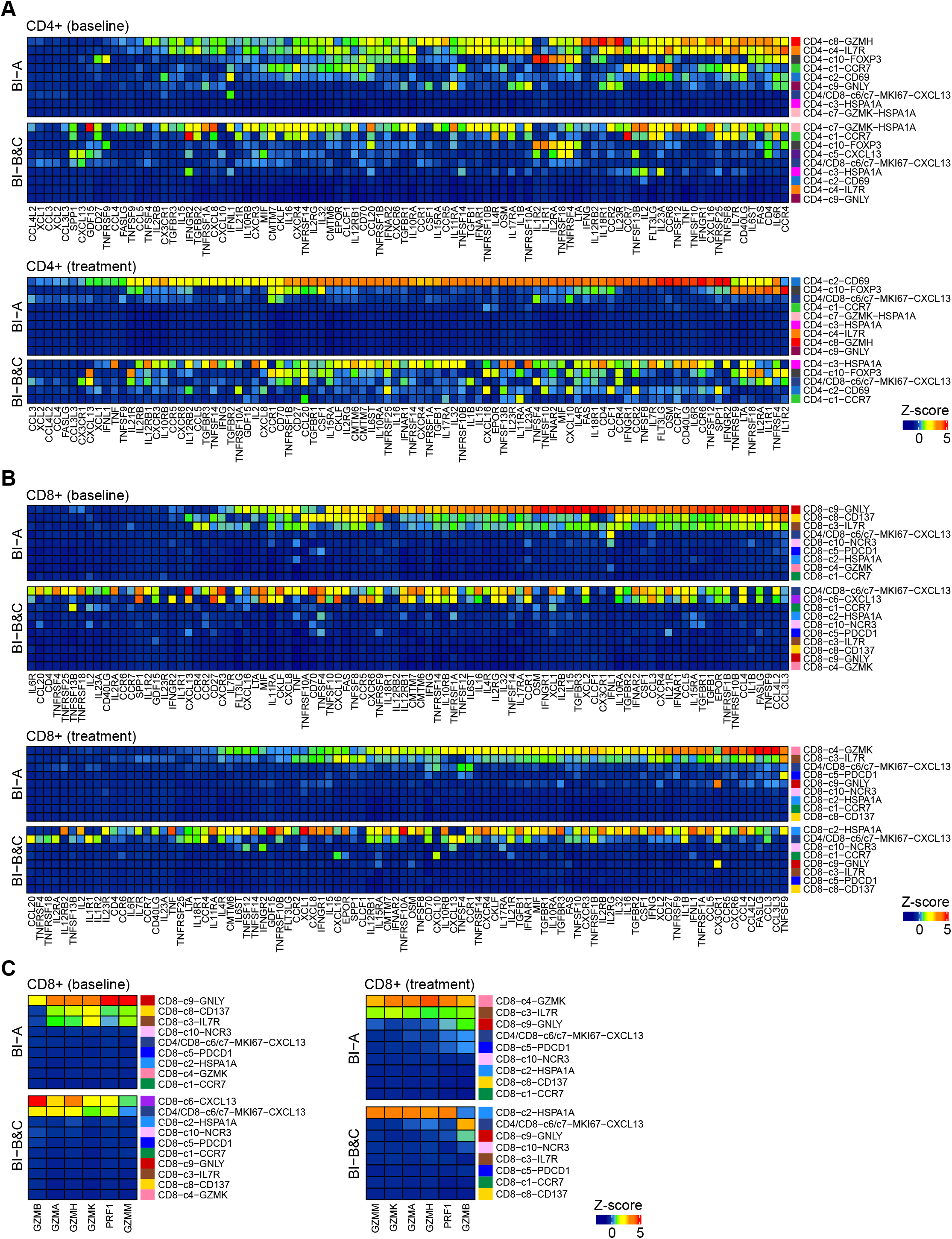
Landscape of cytokines, chemokines, granzymes and perforin produced by T cells in response to treatment. (A) Cytokines and chemokines produced by each CD4^+^ T-cell subtype from BI-A and BI-B&C at baseline (top) or after treatment (bottom). (B) Cytokines and chemokines produced by each CD8^+^ T-cell subtype from BI-A and BI-B&C at baseline (top) or after treatment (bottom). (C) Granzymes and perforin produced by each CD8^+^ T-cell subtype from BI-A and BI-B&C at baseline (left) or after treatment (right). In (A-C), gene expression was weighted by the percentage of each T-cell subtype, followed by z-score transformation.

---

**Table S1.** Clinical information of 37 primary liver cancer patients involved in this study.

| No. | ID<sup>a</sup> | Sex | Age<sup>b</sup> | Race | Diagnosis | Stage | Etiology | Systematic or Surgical Treatment<sup>c</sup> | Mode of Action<sup>d</sup> | Local Therapy<sup>e</sup> |
| --- | --- | --- | --- | --- | --- | --- | --- | --- | --- | --- |
| 1 | H08 | male | 66 | African American | HCC | IV | HCV | Tremel | CTLA-4 | TACE |
| 2 | H18 | male | 61 | Caucasian | HCC | IV | HCV | Durva/Tremel | PD-L1/CTLA-4 | XRT |
| 3 | H21 | male | 77 | Caucasian | HCC | III | None | Resection | NA | NA |
| 4 | H23 | female | 41 | Caucasian | HCC | III | Fatty liver | Durva/Tremel | PD-L1/CTLA-4 | TACE |
| 5 | C25 | female | 47 | Caucasian | iCCA | IV | None | Durva/Tremel | PD-L1/CTLA-4 | NA |
| 6 | C26* | male | 63 | Caucasian | iCCA | IV | None | Durva/Tremel | PD-L1/CTLA-4 | NA |
| 7 | H28 | female | 63 | African American | HCC | III | HCV | Durva/Tremel | PD-L1/CTLA-4 | NA |
| 8 | C29 | male | 61 | Caucasian | iCCA | IV | None | Durva/Tremel | PD-L1/CTLA-4 | NA |
| 9 | H30 | male | 63 | African American | HCC | IV | HCV | Durva/Tremel | PD-L1/CTLA-4 | XRT |
| 10 | H34* | male | 63 | Caucasian | HCC | IV | HCV | Durva/Tremel | PD-L1/CTLA-4 | TACE |
| 11 | C35 | female | 64 | Caucasian | iCCA | IV | None | Pembro | PD-1 | NA |
| 12 | H37 | male | 65 | African American | HCC | IV | HCV | Durva/Tremel | PD-L1/CTLA-4 | NA |
| 13 | H38 | male | 74 | Asian | HCC | I | None | Resection | NA | NA |
| 14 | C39 | male | 61 | Caucasian | iCCA | IV | None | Pembro | PD-1 | NA |
| 15 | H41 | male | 61 | Caucasian | HCC | IV | HCV | No treatment | NA | NA |
| 16 | C42 | male | 67 | Asian | iCCA | IV | None | Pembro | PD-1 | NA |
| 17 | H43 | female | 60 | African American | HCC | IV | HBV | Durva/Tremel | PD-L1/CTLA-4 | NA |
| 18 | C46* | male | 69 | Caucasian | iCCA | IV | None | Pembro | PD-1 | NA |
| 19 | H49* | male | 54 | African American | HCC | IV | HCV | Durva/Tremel | PD-L1/CTLA-4 | TACE |
| 20 | C52 | female | 49 | Caucasian | iCCA | IV | None | Pembro | PD-1 | NA |
| 21 | H54 | male | 35 | African American | HCC | IV | HBV | Durva/Tremel | PD-L1/CTLA-4 | NA |
| 22 | H55 | female | 70 | African American | HCC | IV | None | Durva/Tremel | PD-L1/CTLA-4 | RFA |
| 23 | C56 | female | 52 | Caucasian | iCCA | IV | None | Durva/Tremel | PD-L1/CTLA-4 | RFA |
| 24 | H58* | male | 73 | Hispanic | HCC | IV | None | Durva/Tremel | PD-L1/CTLA-4 | TACE |
| 25 | C60 | female | 80 | Caucasian | iCCA | III | None | Resection | NA | NA |
| 26 | H62 | female | 67 | Caucasian | HCC | I | HCV | Resection | NA | NA |
| 27 | H63 | male | 81 | Asian | HCC | IV | HBV | Durva/Tremel | PD-L1/CTLA-4 | RFA |
| 28 | H65 | female | 62 | Caucasian | HCC | IV | HCV | Durva/Tremel | PD-L1/CTLA-4 | NA |
| 29 | C66 | female | 71 | Caucasian | iCCA | IV | None | Pembro | PD-1 | NA |
| 30 | H68* | male | 78 | Asian | HCC | IV | HBV | Durva/Tremel | PD-L1/CTLA-4 | TACE |
| 31 | H70 | male | 72 | Caucasian | HCC | II | None | Resection | NA | NA |
| 32 | H72 | male | 76 | Caucasian | HCC | IV | Fatty liver | No treatment | NA | NA |
| 33 | H73* | male | 58 | Asian | HCC | IV | HBV,HDV | Durva/Tremel | PD-L1/CTLA-4 | RFA |
| 34 | H74 | male | 67 | Caucasian | HCC | IV | HCV | Durva/Tremel | PD-L1/CTLA-4 | RFA |
| 35 | H75 | female | 74 | Asian | HCC | IIIB | HCV | No treatment | NA | TACE |
| 36 | C76 | female | 74 | Caucasian | iCCA | IV | HCV | Durva/Tremel | PD-L1/CTLA-4 | RFA |
| 37 | H77 | male | 72 | Caucasian | HCC | IIIB | HCV | Durva/Tremel | PD-L1/CTLA-4 | TACE |

<sup>a</sup>A total of 46 tumor biopsies were collected, among which 19 were from our previous study. \* indicates cases with $\geq 2$ biopsies collected from different time points.
<sup>b</sup>Age at time of tissue collection used for this study.
<sup>c</sup>Tremel, tremelimumab; Durva, durvalumab; Pembro, pembrolizumab
<sup>d</sup>PD-L1, programmed death-ligand 1; CTLA-4, cytotoxic T-lymphocyte-associated protein 4; PD-1, programmed cell death protein 1; NA, not applicable
<sup>e</sup>TACE, transarterial chemoembolization; RFA, radiofrequency ablation; XRT, radiation therapy

**Table S2.**
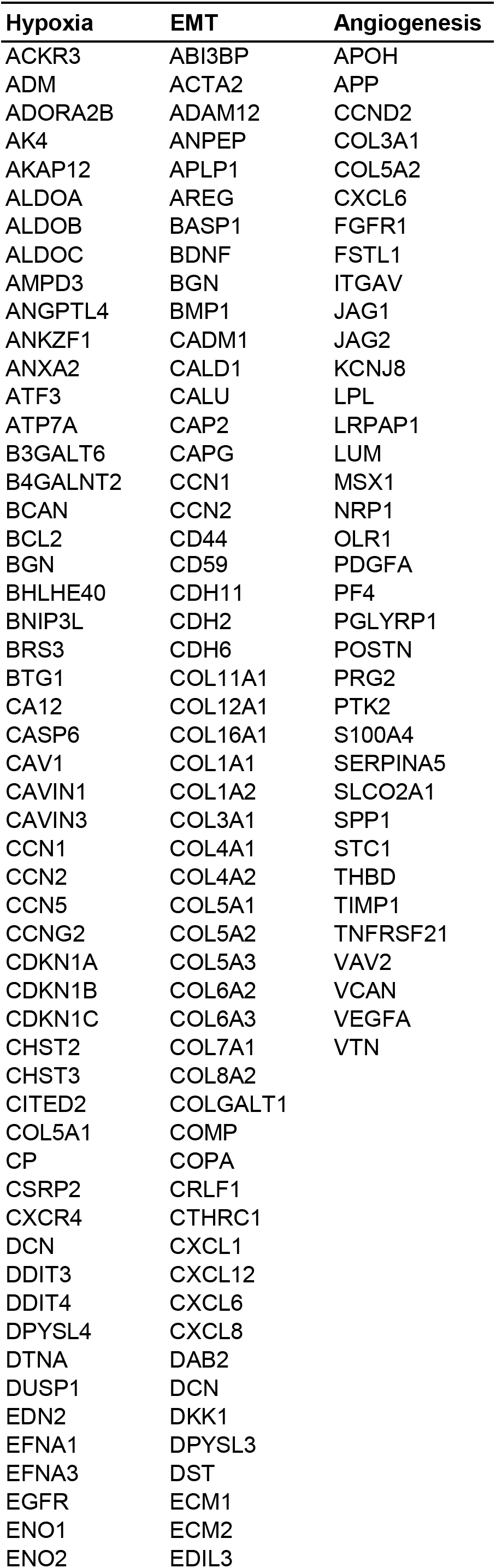
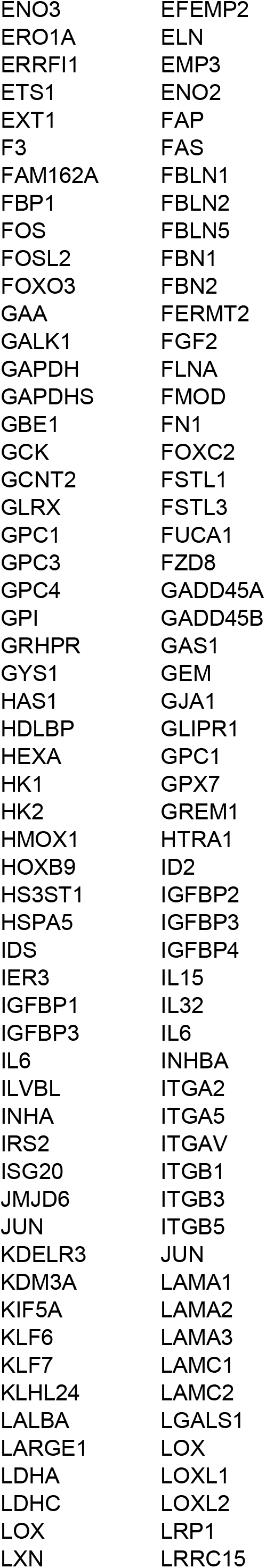
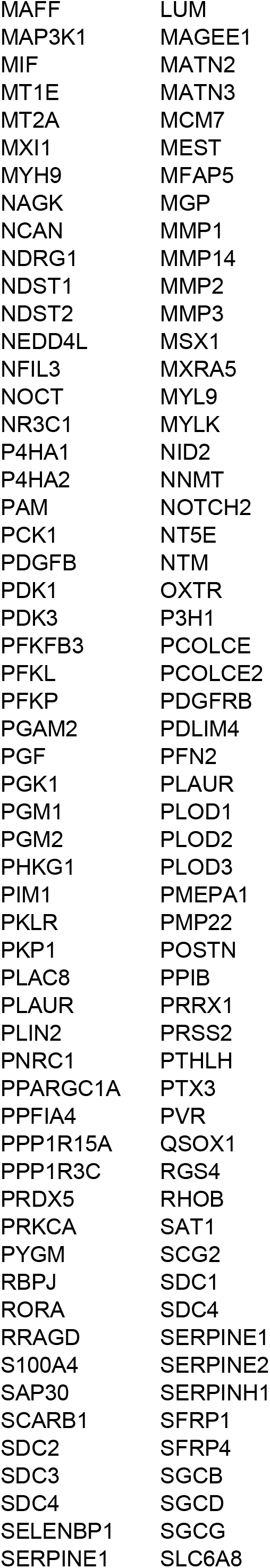
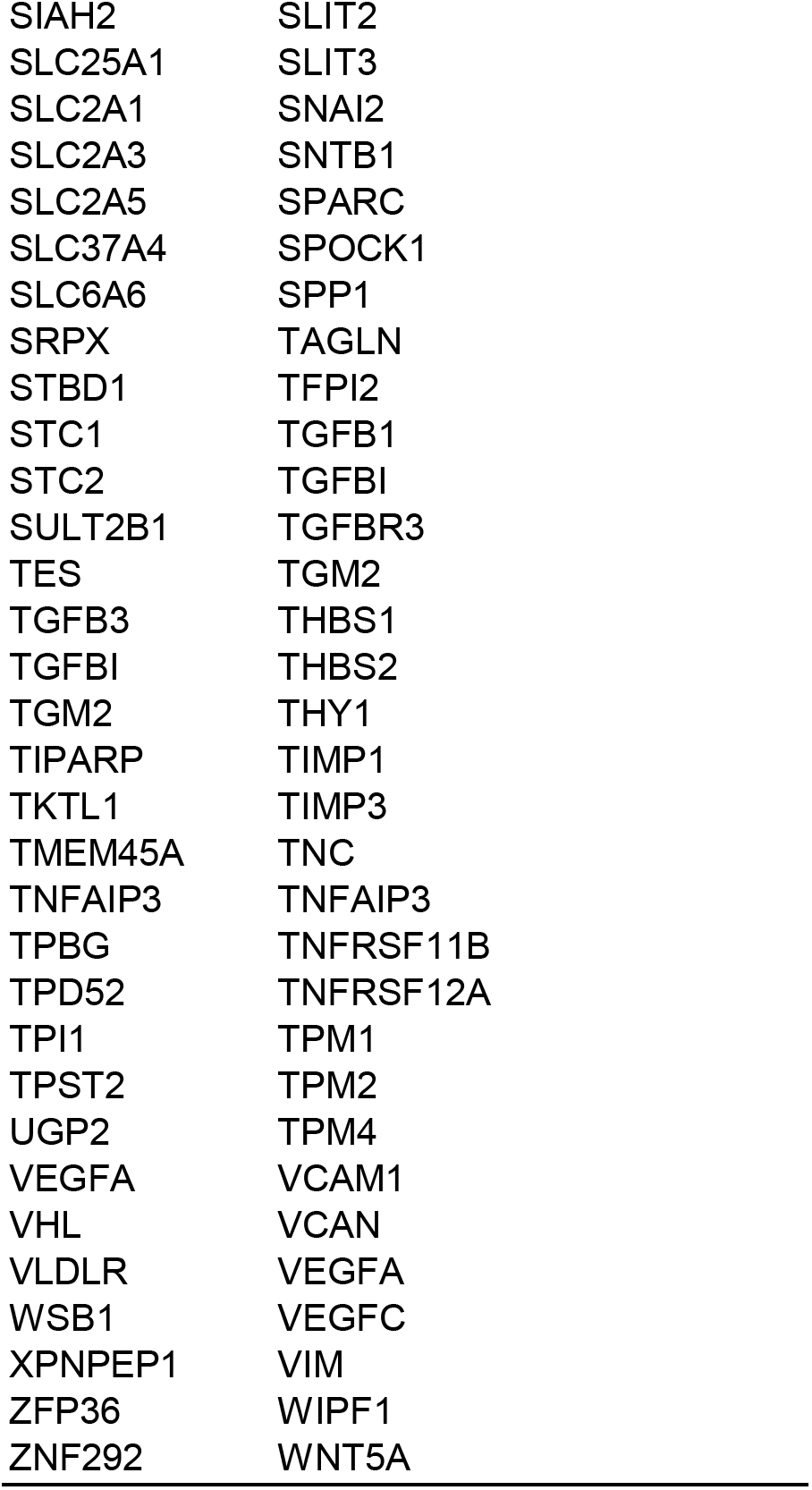
Hypoxia, EMT and angiogenesis related genes.

## REFERENCES

Aizarani, N., Saviano, A., Sagar, Mailly, L., Durand, S., Herman, J.S., Pessaux, P., Baumert, T.F., and Grun, D. (2019). A human liver cell atlas reveals heterogeneity and epithelial progenitors. Nature 572, 199–204.

Alizadeh, A.A., Aranda, V., Bardelli, A., Blanpain, C., Bock, C., Borowski, C., Caldas, C., Califano, A., Doherty, M., Elsner, M., et al. (2015). Toward understanding and exploiting tumor heterogeneity. Nat Med 21, 846–853.

Andor, N., Graham, T.A., Jansen, M., Xia, L.C., Aktipis, C.A., Petritsch, C., Ji, H.P., and Maley, C.C. (2016). Pan-cancer analysis of the extent and consequences of intratumor heterogeneity. Nat Med 22, 105–113.

Azizi, E., Carr, A.J., Plitas, G., Cornish, A.E., Konopacki, C., Prabhakaran, S., Nainys, J., Wu, K., Kiseliovas, V., Setty, M., et al. (2018). Single-Cell Map of Diverse Immune Phenotypes in the Breast Tumor Microenvironment. Cell 174, 1293–1308 e1236.

Boyault, S., Rickman, D.S., de Reynies, A., Balabaud, C., Rebouissou, S., Jeannot, E., Herault, A., Saric, J., Belghiti, J., Franco, D., et al. (2007). Transcriptome classification of HCC is related to gene alterations and to new therapeutic targets. Hepatology 45, 42–52.

Bray, F., Ferlay, J., Soerjomataram, I., Siegel, R.L., Torre, L.A., and Jemal, A. (2018). Global cancer statistics 2018: GLOBOCAN estimates of incidence and mortality worldwide for 36 cancers in 185 countries. CA Cancer J Clin 68, 394–424.

Briones-Orta, M.A., Avendano-Vazquez, S.E., Aparicio-Bautista, D.I., Coombes, J.D., Weber, G.F., and Syn, W.K. (2017). Osteopontin splice variants and polymorphisms in cancer progression and prognosis. Biochim Biophys Acta Rev Cancer 1868, 93–108 A.

Butler, A., Hoffman, P., Smibert, P., Papalexi, E., and Satija, R. (2018). Integrating single-cell transcriptomic data across different conditions, technologies, and species. Nature Biotechnology 36, 411.

Chaisaingmongkol, J., Budhu, A., Dang, H., Rabibhadana, S., Pupacdi, B., Kwon, S.M., Forgues, M., Pomyen, Y., Bhudhisawasdi, V., Lertprasertsuke, N., et al. (2017). Common Molecular Subtypes Among Asian Hepatocellular Carcinoma and Cholangiocarcinoma. Cancer Cell 32, 57–70 e53.

Duffy, A.G., Ulahannan, S.V., Makorova-Rusher, O., Rahma, O., Wedemeyer, H., Pratt, D., Davis, J.L., Hughes, M.S., Heller, T., ElGindi, M., et al. (2017). Tremelimumab in combination with ablation in patients with advanced hepatocellular carcinoma. J Hepatol 66, 545–551.

Finn, R.S., Qin, S., Ikeda, M., Galle, P.R., Ducreux, M., Kim, T.Y., Kudo, M., Breder, V., Merle, P., Kaseb, A.O., et al. (2020). Atezolizumab plus Bevacizumab in Unresectable Hepatocellular Carcinoma. N Engl J Med 382, 1894–1905.

Gerlinger, M., Rowan, A.J., Horswell, S., Larkin, J., Endesfelder, D., Gronroos, E., Martinez, P., Matthews, N., Stewart, A., Tarpey, P., et al. (2012). Intratumor heterogeneity and branched evolution revealed by multiregion sequencing. N Engl J Med 366, 883–892.

Gonzalez-Silva, L., Quevedo, L., and Varela, I. (2020). Tumor Functional Heterogeneity Unraveled by scRNA-seq Technologies. Trends Cancer 6, 13–19.

Gordon, D.M. (2014). The ecology of collective behavior. PLoS Biol 12, e1001805.

Guo, X., Zhang, Y., Zheng, L., Zheng, C., Song, J., Zhang, Q., Kang, B., Liu, Z., Jin, L., Xing, R., et al. (2018). Global characterization of T cells in non-small-cell lung cancer by single-cell sequencing. Nat Med 24, 978–985.

Iacobuzio-Donahue, C.A., Litchfield, K., and Swanton, C. (2020). Intratumor heterogeneity reflects clinical disease course. Nature Cancer 1, 3–6.

Icer, M.A., and Gezmen-Karadag, M. (2018). The multiple functions and mechanisms of osteopontin. Clin Biochem 59, 17–24.

Junttila, M.R., and de Sauvage, F.J. (2013). Influence of tumour micro-environment heterogeneity on therapeutic response. Nature 501, 346–354.

Jusakul, A., Cutcutache, I., Yong, C.H., Lim, J.Q., Huang, M.N., Padmanabhan, N., Nellore, V., Kongpetch, S., Ng, A.W.T., Ng, L.M., et al. (2017). Whole-Genome and Epigenomic Landscapes of Etiologically Distinct Subtypes of Cholangiocarcinoma. Cancer Discov 7, 1116–1135.

Kanayama, M., Xu, S., Danzaki, K., Gibson, J.R., Inoue, M., Gregory, S.G., and Shinohara, M.L. (2017). Skewing of the population balance of lymphoid and myeloid cells by secreted and intracellular osteopontin. Nat Immunol 18, 973–984.

Khatib, S., Pomyen, Y., Dang, H., and Wang, X.W. (2020). Understanding the cause and consequence of tumor heterogeneity. Trends Cancer 6, 267–271.

Kim, D.S., Ryu, J.W., Son, M.Y., Oh, J.H., Chung, K.S., Lee, S., Lee, J.J., Ahn, J.H., Min, J.S., and Ahn, J. (2017). A liver-specific gene expression panel predicts the differentiation status of in vitro hepatocyte models. Hepatology 66, 1662–1674.

Kiselev, V.Y., Kirschner, K., Schaub, M.T., Andrews, T., Yiu, A., Chandra, T., Natarajan, K.N., Reik, W., Barahona, M., and Green, A.R. (2017). SC3: consensus clustering of single-cell RNA-seq data. Nature Methods 14, 483.

Kwon, S.M., Budhu, A., Woo, H.G., Chaisaingmongkol, J., Dang, H., Forgues, M., Harris, C.C., Zhang, G., Auslander, N., Ruppin, E., et al. (2019). Functional genomic complexity defines intratumor heterogeneity and tumor aggressiveness in liver cancer. Scientific Reports 9, 16930.

La Manno, G., Soldatov, R., Zeisel, A., Braun, E., Hochgerner, H., Petukhov, V., Lidschreiber, K., Kastriti, M.E., Lönnerberg, P., and Furlan, A. (2018). RNA velocity of single cells. Nature 560, 494–498.

Lee, J.S., Chu, I.S., Heo, J., Calvisi, D.F., Sun, Z., Roskams, T., Durnez, A., Demetris, A.J., and Thorgeirsson, S.S. (2004). Classification and prediction of survival in hepatocellular carcinoma by gene expression profiling. Hepatology 40, 667–676.

Li, H., Courtois, E.T., Sengupta, D., Tan, Y., Chen, K.H., Goh, J.J.L., Kong, S.L., Chua, C., Hon, L.K., Tan, W.S., et al. (2017). Reference component analysis of single-cell transcriptomes elucidates cellular heterogeneity in human colorectal tumors. Nat Genet 49, 708–718.

Ma, L., Hernandez, M.O., Zhao, Y., Mehta, M., Tran, B., Kelly, M., Rae, Z., Hernandez, J.M., Davis, J.L., Martin, S.P., et al. (2019). Tumor Cell Biodiversity Drives Microenvironmental Reprogramming in Liver Cancer. Cancer Cell 36, 418–430.e416.

Maley, C.C., Aktipis, A., Graham, T.A., Sottoriva, A., Boddy, A.M., Janiszewska, M., Silva, A.S., Gerlinger, M., Yuan, Y., Pienta, K.J., et al. (2017). Classifying the evolutionary and ecological features of neoplasms. Nat Rev Cancer 17, 605–619.

Neftel, C., Laffy, J., Filbin, M.G., Hara, T., Shore, M.E., Rahme, G.J., Richman, A.R., Silverbush, D., Shaw, M.L., and Hebert, C.M.J.C. (2019). An integrative model of cellular states, plasticity, and genetics for glioblastoma. Cell 178, 835–849. e821.

Nowell, P.C. (1976). The clonal evolution of tumor cell populations. Science (New York, NY) 194, 23–28.

Puram, S.V., Tirosh, I., Parikh, A.S., Patel, A.P., Yizhak, K., Gillespie, S., Rodman, C., Luo, C.L., Mroz, E.A., Emerick, K.S., et al. (2017). Single-Cell Transcriptomic Analysis of Primary and Metastatic Tumor Ecosystems in Head and Neck Cancer. Cell 171, 1611–1624 e1624.

Qiu, X., Mao, Q., Tang, Y., Wang, L., Chawla, R., Pliner, H.A., and Trapnell, C. (2017). Reversed graph embedding resolves complex single-cell trajectories. Nature methods 14, 979.

Ramachandran, P., Dobie, R., Wilson-Kanamori, J., Dora, E., Henderson, B., Luu, N., Portman, J., Matchett, K., Brice, M., and Marwick, J. (2019). Resolving the fibrotic niche of human liver cirrhosis at single-cell level. Nature 575, 512–518.

Rittling, S.R., and Chambers, A.F. (2004). Role of osteopontin in tumour progression. BrJ Cancer 90, 1877–1881.

Rittling, S.R., and Singh, R. (2015). Osteopontin in Immune-mediated Diseases. J Dent Res 94, 1638–1645.

Rozenblatt-Rosen, O., Regev, A., Oberdoerffer, P., Nawy, T., Hupalowska, A., Rood, J.E., Ashenberg, O., Cerami, E., Coffey, R.J., Demir, E., et al. (2020). The Human Tumor Atlas Network: Charting Tumor Transitions across Space and Time at Single-Cell Resolution. Cell 181, 236–249.

Salcedo, A., Tarabichi, M., Espiritu, S.M.G., Deshwar, A.G., David, M., Wilson, N.M., Dentro, S., Wintersinger, J.A., Liu, L.Y., Ko, M., et al. (2020). A community effort to create standards for evaluating tumor subclonal reconstruction. Nat Biotechnol 38, 97–107.

Simon, R., Lam, A., Li, M.-C., Ngan, M., Menenzes, S., and Zhao, Y. (2007). Analysis of gene expression data using BRB-array tools. Cancer Informatics 3, 11–17.

Stuart, T., Butler, A., Hoffman, P., Hafemeister, C., Papalexi, E., Mauck III, W.M., Hao, Y., Stoeckius, M., Smibert, P., and Satija, R. (2019). Comprehensive integration of single-cell data. Cell 177, 1888–1902. e1821.

Subramanian, A., Tamayo, P., Mootha, V.K., Mukherjee, S., Ebert, B.L., Gillette, M.A., Paulovich, A., Pomeroy, S.L., Golub, T.R., Lander, E.S., et al. (2005). Gene set enrichment analysis: a knowledge-based approach for interpreting genome-wide expression profiles. Proc Natl Acad Sci USA 102, 15545–15550.

Suzuki, R., and Shimodaira, H. (2006). Pvclust: an R package for assessing the uncertainty in hierarchical clustering. Bioinformatics 22, 1540–1542.

Takafuji, V., Forgues, M., Unsworth, E., Goldsmith, P., and Wang, X.W. (2007). An osteopontin fragment is essential for tumor cell invasion in hepatocellular carcinoma. Oncogene 26, 6361–6371.

Tasic, B., Yao, Z., Graybuck, L.T., Smith, K.A., Nguyen, T.N., Bertagnolli, D., Goldy, J., Garren, E., Economo, M.N., and Viswanathan, S. (2018). Shared and distinct transcriptomic cell types across neocortical areas. Nature 563, 72–78.

TheCancerGenomeAtlasResearchNetwork (2017). Comprehensive and Integrative Genomic Characterization of Hepatocellular Carcinoma. Cell 169, 1327–1341 e1323.

Tirosh, I., Izar, B., Prakadan, S.M., Wadsworth, M.H., 2nd, Treacy, D., Trombetta, J.J., Rotem, A., Rodman, C., Lian, C., Murphy, G., et al. (2016a). Dissecting the multicellular ecosystem of metastatic melanoma by single-cell RNA-seq. Science (New York, NY) 352, 189–196.

Tirosh, I., Venteicher, A.S., Hebert, C., Escalante, L.E., Patel, A.P., Yizhak, K., Fisher, J.M., Rodman, C., Mount, C., Filbin, M.G., et al. (2016b). Single-cell RNA-seq supports a developmental hierarchy in human oligodendroglioma. Nature 539, 309–313.

Trapnell, C., Cacchiarelli, D., Grimsby, J., Pokharel, P., Li, S., Morse, M., Lennon, N.J., Livak, K.J., Mikkelsen, T.S., and Rinn, J.L. (2014). The dynamics and regulators of cell fate decisions are revealed by pseudotemporal ordering of single cells. Nature Biotechnology 32, 381.

Venteicher, A.S., Tirosh, I., Hebert, C., Yizhak, K., Neftel, C., Filbin, M.G., Hovestadt, V., Escalante, L.E., Shaw, M.L., Rodman, C., et al. (2017). Decoupling genetics, lineages, and microenvironment in IDH-mutant gliomas by single-cell RNA-seq. Science (New York, NY) 355.

Vento-Tormo, R., Efremova, M., Botting, R.A., Turco, M.Y., Vento-Tormo, M., Meyer, K.B., Park, J.-E., Stephenson, E., Polański, K., and Goncalves, A. (2018). Single-cell reconstruction of the early maternal– fetal interface in humans. Nature 563, 347.

Waltman, L., and Van Eck, N.J. (2013). A smart local moving algorithm for large-scale modularity-based community detection. The European Physical Journal B 86, 471.

Wang, X.W., and Thorgeirsson, S.S. (2014). The biological and clinical challenge of liver cancer heterogeneity. Hepat Oncol 1, 349–353.

Worns, M.A., and Galle, P.R. (2014). HCC therapies--lessons learned. Nat Rev Gastroenterol Hepatol 11, 447–452.

Ye, Q.H., Qin, L.X., Forgues, M., He, P., Kim, J.W., Peng, A.C., Simon, R., Li, Y., Robles, A.I., Chen, Y., et al. (2003). Predicting hepatitis B virus-positive metastatic hepatocellular carcinomas using gene expression profiling and supervised machine learning. NatMed 9, 416–423.

Zhang, L., Yu, X., Zheng, L., Zhang, Y., Li, Y., Fang, Q., Gao, R., Kang, B., Zhang, Q., Huang, J.Y., et al. (2018). Lineage tracking reveals dynamic relationships of T cells in colorectal cancer. Nature 564, 268–272.

Zhang, Q., He, Y., Luo, N., Patel, S.J., Han, Y., Gao, R., Modak, M., Carotta, S., Haslinger, C., and Kind, D. (2019). Landscape and dynamics of single immune cells in hepatocellular carcinoma. Cell 179, 829–845. e820.

Zheng, C., Zheng, L., Yoo, J.K., Guo, H., Zhang, Y., Guo, X., Kang, B., Hu, R., Huang, J.Y., Zhang, Q., et al. (2017). Landscape of Infiltrating T Cells in Liver Cancer Revealed by Single-Cell Sequencing. Cell 169, 1342–1356 e1316.

Zhong, S., Zhang, S., Fan, X., Wu, Q., Yan, L., Dong, J., Zhang, H., Li, L., Sun, L., Pan, N., et al. (2018). A single-cell RNA-seq survey of the developmental landscape of the human prefrontal cortex. Nature 555, 524–528.

